# Full-Field Analysis Indicates Late Reperfusion Therapy Broadens and Mechanically Smooths the Borderzone During Post-Infarction Inflammation

**DOI:** 10.1101/2025.09.02.673841

**Authors:** Daniel P. Pearce, Tianyi Zheng, Paul Campagnola, Colleen M. Witzenburg

## Abstract

Late reperfusion therapy (LRT; ≥ 3 hours post-MI) significantly reduces the risk of ventricular rupture following myocardial infarction (MI), yet the structural and mechanical mechanisms behind this protection remain unclear. We hypothesized that LRT would alter the biomechanical properties of the infarct borderzone and to investigate this, we utilized laser micrometry, planar biaxial testing, and quantitative polarized light imaging (QPLI) to quantify spatial variations in the geometric, mechanical, and structural properties of the left ventricle extracellular matrix (LV ECM) in adult male Sprague-Dawley rats. Rats received permanent occlusion (PO), LRT, or a sham surgery and tissue was collected 1-day post-MI, during the inflammatory phase of healing. LRT generated a larger infarct borderzone (LRT: 31.5 ± 7.6 mm^2^; PO: 22.5 ± 4.2 mm^2^; *p* < 0.05) in comparison to PO. Infarct core and borderzone stiffness was reduced post-MI, and LRT samples exhibited smoother, more consistent stiffness gradients between infarct core and remote regions than PO samples. In general, infarcted LV ECM from were thicker and more spatially variable than sham samples, but less stiff. Additionally, dynamic QPLI revealed decreased collagen fiber alignment in infarct cores relative to borders, though this did not differ between PO and LRT groups. Complementary second harmonic generation imaging revealed more gradual, consistent transitions in collagen fiber alignment throughout LV ECMs subjected to LRT, although this was limited to one sample from each group. Ultimately, these results further justify LRT and may inform future therapeutic strategies aimed at spatially modulating post-MI tissue mechanics to improve patient outcomes.

## 1. Introduction

Occlusion of the coronary arteries leads to myocardial infarction (MI), irreparable cardiomyocyte damage, and impaired left ventricle (LV) function for nearly a million Americans, and millions more worldwide, each year [1,2]. Following occlusion, an intense inflammatory response occurs in the infarcted myocardium [3–5], during which numerous immune cells, including neutrophils [6–8] and macrophages [9–12] migrate towards the infarcted region. As they infiltrate the necrotic core of the infarction, these cells release pro-inflammatory cytokines and matrix metalloproteinases (MMPs) that degrade the native extracellular matrix (ECM) and contribute to accentuated immune cell infiltration [11,13–19]. The destruction of the native ECM is essential for removing necrotic cardiomyocytes, scar formation, and long-term healing [4,20]. Unfortunately, its degradation also leaves the infarcted myocardium defenseless against wall thinning [21–24], rising LV volume [25], and an elevated risk of ventricular rupture (an often fatal post-MI complication) during the first week after a heart attack [14,26–28].

Reperfusion therapy (RT) is one of the most effective strategies for managing adverse effects of MIs, including ventricular rupture [29–36]. Reperfusion of the coronary vasculature can be achieved pharmacologically through the administration of fibrinolytics [37–40], mechanically through the use of balloon angioplasty or placement of a stent [41–44], or through a combination of these approaches [31,45]. Sadly, the effectiveness of RT depends upon the timeliness of the procedure: the sooner the blocked artery is reopened, the fewer ischemic cardiomyocytes are lost and the less extensive the resulting myocardial damage [29,35,46–48]. Administering RT this early, though, is a challenge for many clinics, especially those treating patients from rural and underserved communities [49–53]. As a result, late reperfusion therapy (LRT; ≥ 3 hours post-MI) is more common in clinical practice. Although LRT fails to salvage ischemic cardiomyocytes and limit infarct size, it has been shown to limit adverse post-MI outcomes [24,54–57]. LRT limits infarct expansion [57], preserves the geometry of the LV wall [24], and reduces rates of ventricular rupture between 3 – 5 days post-MI [14,26–28]. This time frame corresponds to a critical period in patients when the walls of the LV are thinning [21–24], LV volumes are increasing [25], and the local ECM is being dismantled [11,13–19]. However, the basis for the protective effects of LRT are not yet fully understood.

While prior studies present compelling evidence that LRT reduces ventricular rupture post-MI, including complete prevention of rupture in mice [35], there remains no clear consensus on the underlying mechanism responsible for this benefit. Collagen content has widely been proposed as a potential predictor of post-MI ventricular rupture [28,58–60], but findings across a range of animal models do not support this theory [35,61–64]. For example, despite a two-fold increase in rupture incidence, Ma et al. [35] reported no significant difference in collagen content between wild type and MMP null mice at 3- and 5-days post-MI, when rupture risk peaks. In larger animal models (rats, rabbits, and dogs), collagen content, as assessed by hydroxyproline assays, was similar between permanently occluded (PO) and LRT groups 2 to 4 weeks post-MI [62–64]. While outside the inflammatory phase of healing, when the heart is most vulnerable, these studies also measured tissue strength. Connelly et al. [63] reported a two-fold reduction in uniaxial tensile strength in LRT vs. PO samples with strength correlating to collagen crosslinking – not content – measured via NaB^3^H_4_ reduction and an Aldol assay. After experiments in rats and four different strains of mice, Fang et al. likewise determined that collagen cross-linking, quantified by insoluble collagen content, correlated inversely with rupture risk [65,66]. However, in their subsequent study, older mice had twice the rupture incidence as young mice despite having 60% more insoluble collagen [59]. Additionally, none of these studies utilized any form of RT, leaving the relationship between RT, collagen cross-linking and organization, and ventricular rupture risk unresolved [65].

Accelerated healing is another proposed beneficial effect of LRT [56,67–70], potentially reducing rupture incidence by simply shortening the post-MI healing period when the heart is vulnerable. However, semi-quantitative histology scores of collagen content and immune cell infiltration showed similar patterns of healing in LRT and PO groups, despite LRT samples exhibiting greater resistance to *ex vivo* rupture and tearing throughout the first week post-MI in rabbits [69]. These previous studies have largely relied on uniaxial tensile testing (prohibiting analysis of mechanical anisotropy; [63,69]), hydroxyproline assays (providing a one-dimensional estimate of collagen content; [62–64,67,71]), and homogeneous samples taken from the infarct core [62–64,67,69,71]. Full-field characterizations of the heterogeneous LV ECM during post-MI inflammation are rare, if not totally absent, in the literature. Consequently, our understanding of how the infarcted ECM is organized, how its organization varies between different regions of the heart, and how it behaves mechanically under physiologically relevant loading conditions during the first few days post-MI remains incomplete [3].

Despite decades of research, the existence of a mechanically, structurally, and biologically distinct borderzone surrounding the infarct core remains one of the most debated and elusive concepts in the field [3,21,72–77]. Infarct borders have been associated with notable immune cell infiltration [11,13,14,65,78], decreased wall thicknesses and elevated wall stresses [69,79–81], and are a common site of ventricular rupture [14,65,69,82,83]. Their size, mechanical function, and structural organization, however, have been understudied, particularly following LRT. This has resulted in a lack of information regarding the effects of LRT on borderzone properties and subsequent rates of ventricular rupture. In this study, we utilize a combination of novel, full-field approaches (laser micrometry [84], planar biaxial testing [84,85], and quantitative polarized light imaging (QPLI; [86]) to explore the spatially variable biomechanical properties of infarcted LV ECM and the modulatory effects of LRT. These full-field approaches allowed us to investigate our hypothesis that LRT generates a broader border between the infarct core and remote tissue, effectively limiting mechanical and structural heterogeneity throughout the heart, reducing stress concentrations at the edges of the infarct core, and creating a smoother transition in properties between the infarct core and remote tissue, as opposed to PO. Our findings emphasize several therapeutic advantages of LRT, which we hope reinforce the significance of LRT for MI management and encourage novel spatially conscious treatment strategies for improving MI remodeling as healing progresses and heart failure becomes a dominant risk factor [87–89].

## 2. Methods

To investigate post-MI remodeling and the beneficial effects of LRT, we employed a combination of full-field approaches to produce comprehensive descriptions of the mechanical and structural properties of decellularized LV ECM. We first detail our experimental approaches, including the animal model and whole-organ perfusion decellularization, as well as sample preparation and our laser micrometry protocol, through which we obtained full-field thickness profiles for each sample. We then describe our QPLI and mechanical testing protocols, which allowed us to generate full-field structural and mechanical characterizations for each sample. Finally, we describe our workflow for data processing and analysis, including the application of a *k*-means clustering algorithm to produce regional descriptions of tissue structure and mechanics.

### 2.1 Experimental Approaches

#### 2.1.1 Animal Model of Myocardial Infarction

Adult male Sprague Dawley rats (weight: 492.7 ± 18.1 g) were randomly assigned to PO via ligation of the left anterior coronary artery; late reperfusion therapy (LRT), in which the left anterior coronary artery was ligated and then released after three hours of ischemia; or a sham surgery, in which each rodent’s chest was opened surgically but their hearts were unperturbed. Using a non-parametric Kruskal-Wallis test, no differences in pre-surgery rodent weight were detected between groups (sham: 472.5 ± 47.2 g; PO: 498.2 ± 69.2 g; LRT: 507.5 ± 35.3 g; Figure S1). MI was confirmed visually by the gross blanching of myocardium downstream of the suture. Hearts were harvested one (1) day after the initial surgery, flushed with a solution of 2,3-butanedione monoxime (BDM; Sigma-Aldrich) and 1% PBS (Sigma-Aldrich), and then stored in this solution at -80 degrees Celsius. They were removed from the freezer and thawed for 24 hours at 3 degrees Celsius before decellularization and subsequent mechanical testing. For this study, a total of 31 animals underwent surgery (sham: 12 rodents; PO: 9 rodents; LRT: 10 rodents). All animal housing and surgeries, managed by the University of Wisconsin-Madison School of Medicine and Public Health Biomedical Research Model Services team, were conducted according to institutional and NIH guidelines.

#### 2.1.2 Whole-Organ Decellularization

Each heart was decellularized by retrograde perfusion of sodium dodecyl sulfate (SDS; Sigma-Aldrich) and Triton X-100 (Sigma-Aldrich; [90]), a procedure shown to have negligible effects on the stiffness and mechanical alignment of rodent right ventricle [91]. Briefly, the heart was perfused with SDS via constant pressure (gravity driven flow at ∼7.5 kPa/56 mmHg) for approximately 24 hours, then constant flow (2 mL/min; Masterflex L/S Digital Precision Modular Drive) for 24 hours or until the heart became visibly pale, slightly transparent, and decellularized. Hearts were then perfused with deionized water and Triton X-100 for 30 minutes each at constant pressure, then rinsed with PBS (constant flow at 2 mL/min) for 48 – 72 hours. Isolation of the ECM resulted in thinning of the LV wall, enabling plane stress characterizations of the LV’s passive mechanical properties (i.e., length:thickness ≥ 10; [92,93]). Four hearts were excluded from additional testing or analysis due to damage during the decellularization process (1 sham, accidental aspiration into the decellularization system) or incomplete decellularization (1 PO and 2 LRT, visible red/pink regions). Histology (H&E, Picrosirius Red, and Alcian blue; Figures S2 – S4, respectively) revealed the absence of residual cardiomyocytes and no consistent differences in cellularity between groups or in various regions of the remaining hearts, leading us to conclude that decellularization had similar efficacy on sham, PO, and LRT hearts.

#### 2.1.3 Sample Preparation and Laser Micrometry

For each decellularized heart (Figure 1A, B), the atria and right ventricle were removed (Figure 1C). Next, the LV was cut longitudinally down the interventricular septum, resulting in an oval-shaped sample that included the entire LV free wall and portions of the septum on both circumferential edges (Figure 1D). Using biopsy punches and razor blades, the LV free wall was trimmed to a cruciform shape such that the circumferential and longitudinal axes of the heart were aligned with the horizontal and vertical arms of the sample, respectively (Figure 1E, F; [94,95]). Each PO and LRT sample were formed to include both infarcted and remote LV. Sham samples included as much of the LV free wall as possible. Each sample was then scanned by a LJ-V7080 class II laser micrometer (Keyence; Itasca, IL) operating at a scanning frequency of 200 Hz to capture full-field thickness [84,86]. A custom MATLAB script (available for download from Pearce et al. [84]) generated 3D thickness profiles from the raw CSV output files.

**Figure 1.**
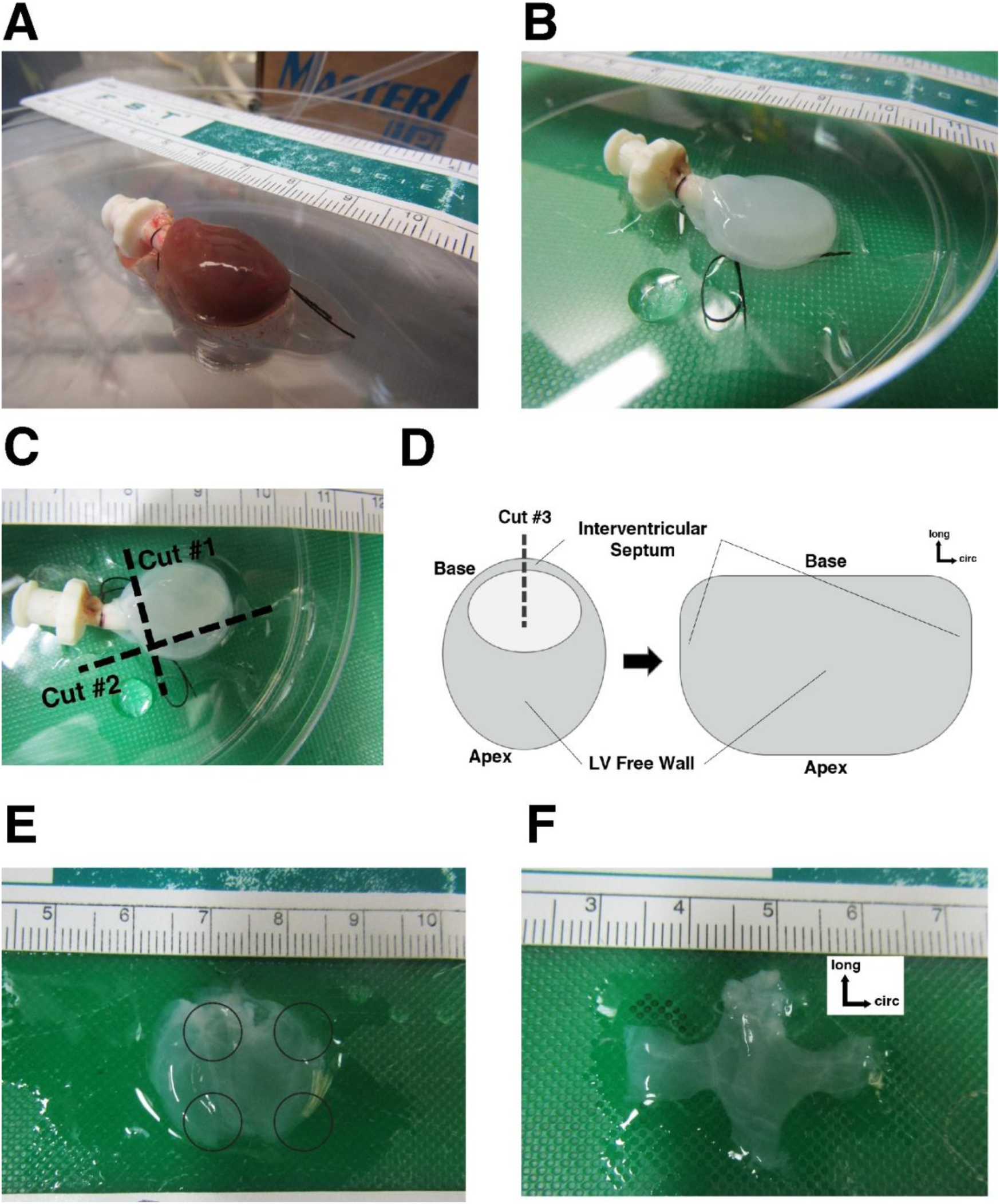
A representative sham rat heart prior to (A) and following (B) whole-organ perfusion decellularization. (C) Cut #1 removed the atria, connective tissue, and luer lock from the decellularized heart and cut #2 removed the right ventricle. (D) Cut #3 was made down the interventricular septum to create a planar, oval-shaped slab of decellularized left ventricle with the free wall in the center. The free wall of the representative sample with locations (black circles) of 5-mm biopsy punches (E) used to form a cruciform shape (F) for planar biaxial testing.

#### 2.1.4 Planar Biaxial Testing: Dynamic QPLI

Next, we conducted dynamic QPLI, a novel technology which capitalizes on the birefringent properties of collagen fibers to quantify the strength and direction of fiber alignment in thin collagenous tissues [86,96–99], during planar biaxial testing. Samples were secured in our biaxial testing machine (TestResources; Shakopee, MN, USA) using a custom clamping system and immersed in PBS [84]. The QPLI lens and camera (FLIR Blackfly BFS-U3-51S5P-C; Wilsonville, OR, USA) collected images of the sample surface at a frequency of 7 Hz [86]. The resolution of these images was 122.1 ± 2.3 px/mm for the sham samples, 114.2 ± 9.7 px/mm for the PO samples, and 118.8 ± 4.5 px/mm for the LRT samples. Imaging resolution did not differ between groups (Figure S5). All samples were preloaded to ≤ 15 kPa, which was less than 10% of the maximum load expected during testing [100]. Samples were then preconditioned using ten sequential equibiaxial extensions (λ_x_ = λ_y_ = 1.2) to achieve a state of pseudoelasticity characterized by repeatable force-displacement curves [84,100]. Then, they were subjected to our unique planar biaxial testing protocol, which was intentionally designed to generate a broad and heterogeneous strain space [84,85,95,101] and involves fifteen different biaxial extensions (including asymmetric single-arm, adjacent-arm, and three-arm extensions, as well as more conventional strip uniaxial and equibiaxial extensions). Comprehensive details of the mechanical testing protocol are available in Pearce et al. [84] The Stokes parameters (*S0*, *S1*, and *S2*) were computed from the raw intensity of transmitted incident light at various angles captured by “superpixels” in the camera’s division-of-focal plane sensor [97,99], defined as:

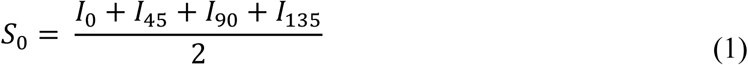

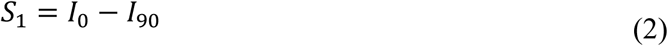

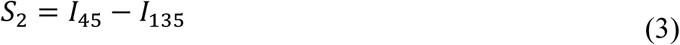

where *I_j_* indicates the incident light at angle *j* in degrees measured by each superpixel. Then the Stokes parameters were used to calculate two key metrics, the degree of linear polarization (*DoLP*) and the angle of polarization (*AoP*), defined as:

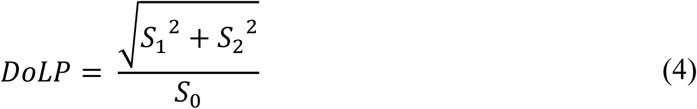

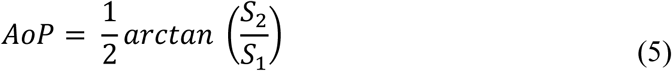

where AoP is on a circular scale from 0° to 180° and indicates the average collagen fiber orientation within a pixel. DoLP, the degree of linear polarization, ranges from 0 (no polarization) to 1 (total polarization) and is a measure of the strength of collagen fiber alignment within a pixel. When attaching each sample to our mechanical testing system, we aligned the circumferential axis of the LV with the horizontal axis of both the testing system and the QPLI camera (i.e., AoP = 0°). As aforementioned, cruciform samples were prepared so that the circumferential and longitudinal directions of each sample were aligned with actuator arms and mechanical testing axes (Figure 1). For more information regarding the development of QPLI and the influence of a sample’s structure and geometry on QPLI outputs, we refer the reader to work by York et al. [97], King et al. [99], and Iannucci et al. [98]. The details of our QPLI protocol are available in Sreedhar et al. [86].

#### 2.1.5 Planar Biaxial Testing: Digital Image Correlation and Force Acquisition

We then applied random speckle patterns to each sample and repeated all fifteen biaxial extensions to measure dynamic full-field Green-Lagrange strain and boundary forces. We drained the tissue bath such that the sample was no longer submerged, manually added a charcoal speckle pattern to the sample’s surface, and allowed the pattern to dry for ∼10 minutes. We then refilled the tissue bath with PBS, removed the QPLI lens and camera from the overhead camera mount, and replaced it with a standard, non-polarized camera system (Imperx, PoE-C2400 Camera, 2464 × 2056 pixels, 5 megapixels, 36 fps, Boca Raton, FL, USA; Computar M3Z1228C-MP Lens, Mebane, NC, USA) to collect images for strain tracking [84]. The resolution of these images was 114.8 ± 26.1 px/mm for the sham samples, 95.6 ± 18.3 px/mm for the PO samples, and 97.5 ± 7.7 px/mm for the LRT samples and did not differ between groups (Figure S6). Each sample was connected to a combination of one degree-of-freedom (1DOF; TestResources, WF12S Miniature Fatigue Resistant Submersible IP65 Load Cells, 0.006 N resolution) and 6DOF (ATI, Nano 17 IP68 F/T Transducers, 0.006 N resolution; Apex, NC, USA) load cells, which measured both boundary normal and shear forces throughout mechanical testing. Upon completion of this protocol, a final equibiaxial extension was performed to evaluate if the sample maintained its pseudoelastic state or sustained damage during mechanical testing. Seven samples (3 sham, 2 PO, and 2 LRT) tore or were damaged (> 10% drop in peak force between the first and final equibiaxial extensions) during mechanical testing and were excluded from further analysis. One additional sham sample was excluded due to equipment malfunction during testing.

### 2.2 Data Analysis

#### 2.2.1 Full-Field Strain Tracking and QPLI Mapping

We utilized custom digital image correlation (DIC) software intentionally designed for heterogeneous soft tissues (available for download from Raghupathy & Barocas [102]) to compute the Green-Lagrange strain in each sample for each extension. First, the image of the sample following speckling, but before the start of the second series of mechanical loading, deemed the undeformed configuration, was meshed with linear Q4 elements (ABAQUS 2018, Dassault Systemes). Then, raw images from all 15 biaxial extensions [84] were compiled and input into the DIC code which computed full-field displacements and Green-Lagrange strain tensors.

By maintaining the same unloaded configuration during dynamic QPLI and standard imaging of speckled samples, we were able to map QPLI results to the DIC mesh. The locations of the grips and sample boundaries in the undeformed image were identified for the DIC mesh and QPLI. Once this mapping was established, and as the DIC mesh deformed throughout different extensions, local QPLI metrics were averaged for all pixels within an element. In this way, successful mapping allowed for determination of QPLI metrics in a particular region of the tissue as that region deformed – i.e., capturing the change in DoLP for an element near the top boundary throughout the test as that particular element moved upwards.

#### 2.2.2 Orthotropic Generalized Anisotropic Inverse Mechanics

Our orthotropic Generalized Anisotropic Inverse Mechanics (GAIM) method leverages full-field strain and normal and shear boundary forces to quantify the spatially variable stiffness of heterogeneous soft tissues [85]. Briefly, the GAIM method assumes a linear relationship between the second Piola-Kirchhoff stress tensor and the Green-Lagrange strain tensor to estimate a fourth-order generalization of the linear elasticity tensor, often referred to as the St. Venant stiffness tensor, for each sample partition [84,85,91]. Constraining this model to orthotropy reduces the number of materials constants from six (*K_1111_*, *K_1122_*, *K_1112_*, *K_2222_*, *K_2212_*, and *K_1212_*) to four (*E_1_*, *E_2_*, *μ_12_*, and *φ*) in each region, increasing parameter identifiability and ensuring positive definite, physiologically relevant mechanical characterizations [85]. The measured full-field thickness profiles from laser micrometry were also incorporated into this formulation, enabling GAIM to compute full-field, three-dimensional stiffness tensors (in kPa; [84,85,103]). The eigenvalues of each local stiffness tensor quantify that region’s principal states of stress and strain and the largest eigenvalue, referred to as the first Kelvin modulus (K1), indicates stiffness in the stiffest direction [85,101,104].

The orthotropic GAIM method is nonlinear in its treatment of kinematics, but linear in terms of kinetics, essentially producing a secant stiffness metric. To address this limitation, we applied orthotropic GAIM at three levels of the prescribed boundary strain: low (sham: 6.33 ± 0.31%; PO: 6.45 ± 0.45%; LRT: 6.15 ± 0.0%), medium (sham: 13.7 ± 0.31%; PO: 13.6 ± 0.45%; LRT: 13.9 ± 0.0%), and maximum (all groups = 22%). We computed the percent change in these full-field stiffness metrics between the low-medium strain levels and between medium-maximum strain levels to evaluate nonlinearity in kinetics.

#### 2.2.3 Sample Partitioning

To conduct spatial comparisons of infarcted samples (PO and LRT), we partitioned them into infarct core, border, and remote regions using a *k*-means clustering algorithm [105,106]. This algorithm was applied to full-field S0 (a measure of transmitted light intensity) in the undeformed configuration and each sample was iteratively partitioned into four (1 = infarct core; 2 = border; 3, 4 = remote) distinct regions. To make inter-sample comparisons, we normalized each sample’s infarct core and border values to the mean of the remote region. We also calculated the border-to-core ratio of each metric.

#### 2.2.4 Statistical Analysis

Full-field geometric (thickness scans), mechanical (GAIM), and structural (QPLI datasets) properties were used to produce descriptive statistics including means, standard deviations, and coefficients of variation (CoVs), a metric useful for quantifying spatial heterogeneity [84,86,91,107]. These values were then compared between sham, PO, and LRT groups. Given the small sample sizes in the present study, as well as the non-normal distributions of various full-field measurements, we chose more conservative non-parametric statistical tests to detect statistical differences between groups. To compare sham, PO, and LRT groups, we used a Kruskal-Wallis test and, if statistical differences were detected, *post-hoc* Dunn’s multiple comparison tests. An outlier test employing the Grubbs’ method to compare the full-field stiffness distributions indicated a sham sample was an outlier. Therefore, that sample was excluded. In total, six samples per group (sham *n* = 6; PO *n* = 6; LRT *n* = 6) were included in the final analysis.

For regional comparisons between the PO and LRT groups, we used Mann-Whitney *U*-tests. To compare paired PO values in the infarct core and border, as well as paired LRT values in the infarct core and border, we used a Wilcoxon matched-pairs signed rank test. We then conducted Wilcoxon signed-rank tests to compare infarct and border values to remote values, which had been normalized to a value of one for each sample. All statistical analyses were conducted in GraphPad Prism 10 (GraphPad Software; Boston, MA) and MATLAB R2024b (Mathworks; Natick, MA). A *p*-value < 0.05 was considered statistically significant and values between 0.05 and 0.10 were considered nearing significance.

## 3. Results

LRT has been shown to reduce the risk of ventricular rupture following MI, yet the mechanical and structural mechanisms underlying this cardioprotective effect are poorly understood. We hypothesized that the application of LRT generates a broader border between the infarct core and remote LV, creating a subtler transition in stiffness between these two regions, and reducing sharp mechanical and structural discontinuities. To evaluate this hypothesis, we applied a series of novel, full-field techniques quantifying the alterations in ECM mechanics and structure associated with PO and LRT. First, we quantified the infarct core and border (or, ‘borderzone’) area and width in both the unloaded and loaded states. We then compared full-field and regional thickness profiles of sham, PO, and LRT samples. Building on this, we applied our orthotropic GAIM method[85] to the boundary force and full-field displacement measurements to estimate each sample’s full-field and regional stiffness. We then considered the full-field and regional Green-Lagrange strain distributions during equibiaxial and Strip X extensions. Finally, we compared the full-field and regional DoLP during equibiaxial and Strip X extensions.

### 3.1 LRT Creates a Larger Borderzone

LRT reduces LV rupture incidence post-MI [14,26–28], but the mechanisms behind this benefit are poorly understood. In our study, we employed a *k*-means clustering algorithm to partition each infarcted sample into its infarct core, border, and remote regions (Figure 2A, B) based on full-field transmitted light intensity in the unloaded configuration. As expected, there was no significant difference between the PO and LRT groups when comparing infarct core area (Figure 2C; PO: 33.4 ± 8.9; LRT: 28.9 ± 9.3 mm^2^) or when comparing the infarct core-to-total sample area ratio (Figure S7F; PO: 0.364 ± 0.11; LRT: 0.310. ± 0.11), according to a non-parametric Mann–Whitney *U*-test. Similarly, the total infarct area (infarct core + border relative to the total sample area) was not significantly different between groups (Figure 2E; PO: 0.608 ± 0.12; LRT: 0.644. ± 0.17). These findings suggest that LRT does not alter overall infarct size. However, differences emerged in the borderzone. The LRT group exhibited larger borderzone areas compared to the PO group that neared statistical significance (Figure 2D; PO: 22.5 ± 4.2 mm^2^; LRT: 31.5 ± 7.6 mm^2^, *p* < 0.10), and the ratio of the border-to-core area was significantly higher in the LRT samples (Figure 2F; PO: 0.735 ± 0.30; LRT: 1.15 ± 0.31; *p* < 0.05). There were no significant differences in the remote area or total sample area between the PO and LRT groups (Figure S7C, D).

**Figure 2.**
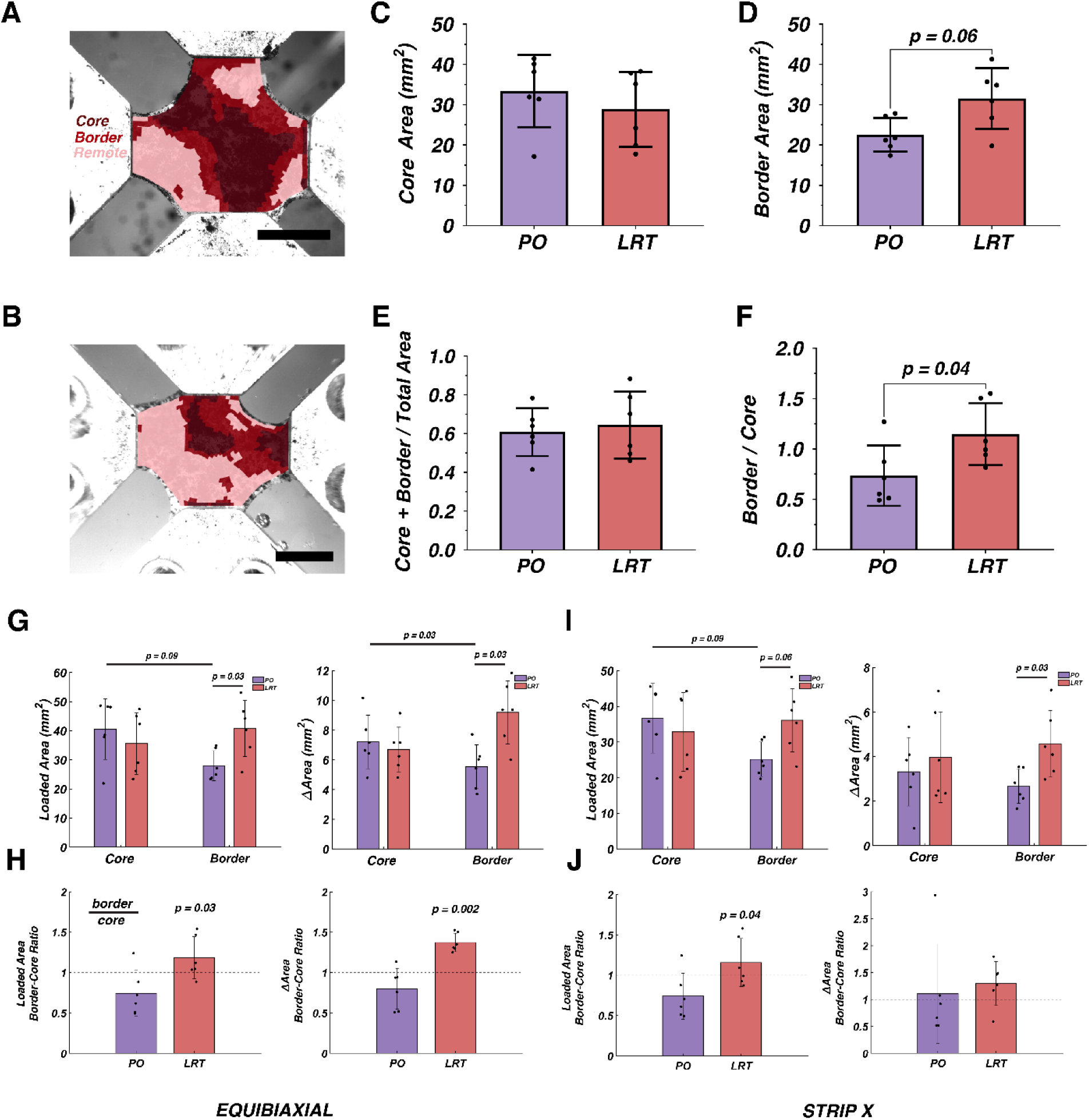
Infarct core, border, and remote regions as identified by a *k*-means clustering algorithm in representative PO (A) and LRT (B) samples. Infarct core (C) and border areas (D) for all PO (*n* = 6) and LRT (*n* = 6) samples, mean ± SD. The ratio of infarct core to total sample area (E) and the border-to-core area ratio (F) for all PO and LRT samples. The peak and change in infarct core and border areas during the equibiaxial extension for all PO and LRT samples (G). The peak and change in the border-to-core area ratio during the equibiaxial extension (H). The peak and change in infarct core and border areas during the Strip X extension for all PO and LRT samples (I). The peak and change in the border-to-core area ratio during the Strip X extension (J). Intra-group statistical significance shown for a Wilcoxon matched-pairs signed rank test and inter-group statistical significance shown for a Mann-Whitney *U*-test. Scale bars indicate 5 mm.

To better quantify the dynamic behavior of the infarcted, decellularized LV ECM, we also measured regional areas in PO and LRT samples during mechanical loading. During equibiaxial and Strip X extensions, infarct cores in PO and LRT samples exhibited comparable areas in the fully loaded state (Figure 2G, I; left) and comparable changes in area with loading (Figure 2G, I; right). Again, differences emerged in the borderzone. In general, the LRT group exhibited a larger loaded borderzone area compared to the PO group (with *p-*values ranging from 0.03 to 0.09 depending on extension, only equibiaxial and Strip X extension data shown; Table S1 includes results for all extensions). Furthermore, the change in border area during each extension was also larger in the LRT group (with *p-*values ranging from 0.009 to 0.24 depending on extension; Table S1). These differences prompted us to compute the loaded border-to-core area ratio and the change in this area ratio as well (Figure 2H, J; Table S2). In the equibiaxial extension, LRT samples exhibited significantly greater border-to-core ratios in both loaded area (∼45% increase; PO: 0.745 ± 0.28; LRT: 1.18 ± 0.26; *p* < 0.05) and change in area (∼53% increase; PO: 0.799 ± 0.25; LRT: 1.37 ± 0.12; *p* < 0.01). In the Strip X extension, LRT samples again exhibited a significantly larger border-to-core loaded area ratio (∼44% increase; PO: 0.740 ± 0.28; LRT: 1.16 ± 0.30; *p* < 0.05); however, the change in area throughout this extension did not differ significantly between groups.

Next, we considered border width. First, we identified peripheral core elements at the infarct core– border interface. For each sample, we then calculated the distance from each peripheral core element to its nearest neighboring remote element, producing an estimate of borderzone width across the entire infarct perimeter (Figure 3A). A non-parametric Mann–Whitney *U*-test indicated no statistical difference in the unloaded borderzone width between the PO and LRT groups (Figure 3B; PO: 1.14 ± 0.29 mm; LRT: 1.38 ± 0.68 mm). Similarly, the loaded borderzone width during equibiaxial extension did not differ between groups (Figure 3C; PO: 1.27 ± 0.30 mm; LRT: 1.56 ± 0.10 mm). This was true for the other extensions as well (with *p*-values ranging from 0.59 to 0.82; Table S3). However, the deformation-induced change in borderzone width (Figure 3D; loaded – unloaded width) was significantly greater in the LRT samples (PO: 0.125 ± 0.02 mm; LRT: 0.187 ± 0.10 mm; *p* < 0.05) during the equibiaxial extension. For all extensions, *p*-values ranged from 0.03 to 0.48, but *p*-values were < 0.10 for only three extensions: the equibiaxial, top and right adjacent arm, and Strip Y extensions (Table S3).

**Figure 3.**
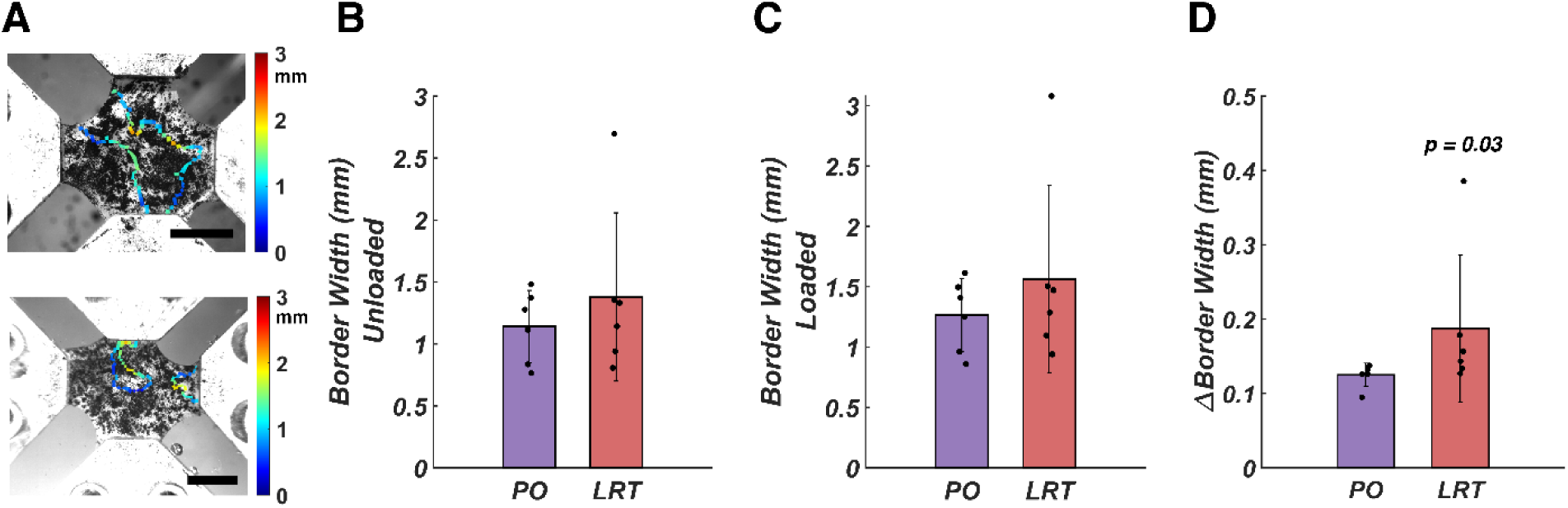
The borderzone width (mm), defined as the distance between each infarct core peripheral element and the nearest healthy remote tissue element for the representative PO and LRT samples (A) shown in Figure 2. The unloaded border width (B) for all PO (*n* = 6) and LRT (*n* = 6) samples, mean ± SD. The peak border width (C) and the change in border width (D) during the equibiaxial extension for all PO and LRT samples. Intra-group statistical significance shown for a Wilcoxon matched-pairs signed rank test and inter-group statistical significance shown for a Mann-Whitney *U*-test. Scale bars indicate 5 mm.

### 3.2 Infarction Increases Thickness and Exacerbates Geometric Heterogeneity

Using a laser micrometer [84], we measured full-field thickness profiles for each sample (Figure 4A, B). Means were calculated from these full-field distributions and compared between sham, PO, and LRT groups using a non-parametric Kruskal-Wallis test and *post-hoc* Dunn’s tests (Figure 4C). Average thicknesses for both PO (1.78 ± 0.31 mm; *p* < 0.05) and LRT samples (1.74 ± 0.33; *p* < 0.05) were significantly greater than sham samples (1.04 ± 0.26 mm). PO and LRT samples also exhibited increased thickness heterogeneity, as quantified by the CoV (Figure 4D; PO: 0.241 ± 0.05; LRT: 0.223 ± 0.06). This difference was only statistically significant for the PO group relative to the sham group (0.148 ± 0.06; *p* < 0.05).

**Figure 4.**
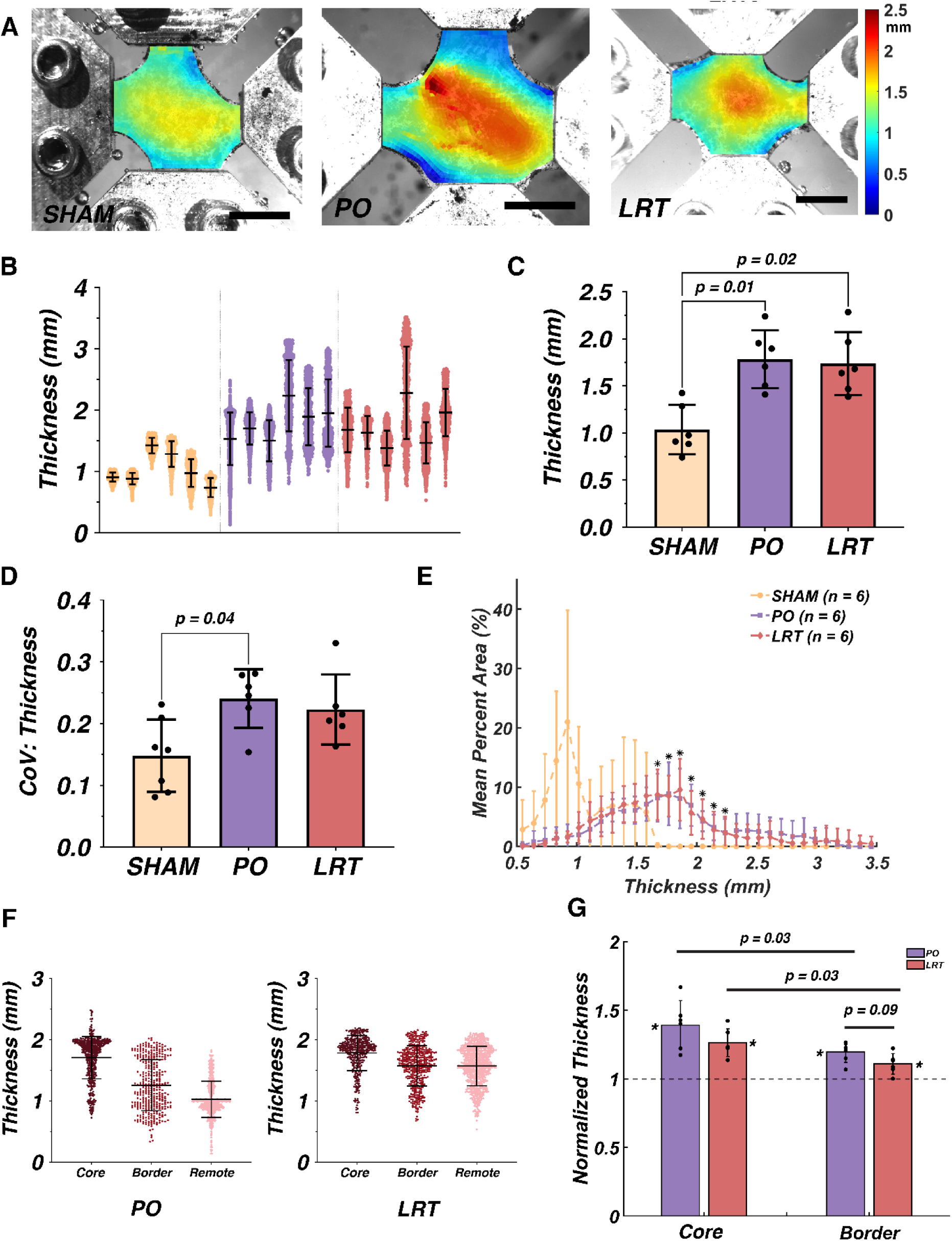
Full-field thickness profiles (A) for representative sham (left), PO (center), and LRT samples (right). Scatter plots showing full-field thickness distributions for each sample (B), a bar plot comparing mean thicknesses (C), and a bar plot comparing the coefficients of variation (CoVs) in thickness (D). (E) The mean percentage of surface area at each thickness value for the sham, PO, and LRT samples. Inter-group statistical significance was detected via a Kruskal-Wallis testing with *post-hoc* Dunn’s tests. (F) Regional partitioning of full-field thickness for the representative samples from Figure 2 with thicknesses for the PO sample on the left and the LRT sample on the right. (G) The infarct core and border mean thickness normalized to the mean peripheral thickness for all PO and LRT samples. In this case, inter-group statistical significance was detected via a Mann-Whitney *U*-test since there were only two groups. An asterisk (*) denotes *p* < 0.05 when regional values were compared to a remote value of one using a Wilcoxon signed-rank test. An octothorpe (#) denotes *p* < 0.10 for a Wilcoxon signed-rank test. Scale bars indicate 5 mm.

To better visualize spatial trends in thickness within each group, we created histograms showing the percent surface area with a particular thickness for each sample (see Figure S8 for representative histograms; [86]). Figure 4E shows the mean percent area for each thickness across sample groups. Sham samples tended to have the most surface area in the 0.5 – 1.5 mm thickness range (∼98% of total surface area), while PO and LRT samples had the most surface area in the 1.5 – 2.5 mm range (∼55% of total surface area). Using a Kruskal-Wallis test and *post-hoc* Dunn’s tests, we found significant differences (*p* < 0.05) between sham and both PO and LRT groups across all bins between 1.67 – 2.23 mm, with no significant differences between PO and LRT groups.

To explore regional trends, we utilized the partitioning scheme previously determined via the *k*-means clustering algorithm. Figure 4F shows the full-field thickness measures partitioned into the infarct core, border, and remote regions of the representative PO and LRT samples shown in Figure 2A and Figure 2B, respectively. In general, the infarct core was thicker than the borderzone in both PO and LRT samples. When considering all the PO and LRT samples, we normalized the core and border thicknesses to peripheral remote values (Figure 4G). The normalized infarct core thicknesses were significantly larger than the neighboring borders: PO: 1.39 ± 0.10 (core) vs. 1.20 ± 0.10 (border); LRT: 1.26 ± 0.08 (core) vs. 1.11 ± 0.07 (border); both *p* < 0.05. While normalized infarct core thicknesses were similar between PO and LRT groups, normalized LRT border thicknesses were slightly decreased and closer to remote values when compared to those in the PO group (*p* < 0.10). It is worth noting, though, that Wilcoxon signed-rank tests indicated both infarct core and borderzone values were significantly larger than one (all *p* < 0.05), their corresponding normalized remote value, for both the PO and LRT groups.

### 3.3 LRT Promotes a Gradual Increase in Stiffness Between Infarct Core and Remote ECM

To quantify spatial stiffness distributions in each sample, we employed the orthotropic GAIM technique, an inverse method that iteratively optimizes the St. Venant stiffness tensor within each region of a sample. This method relies on experimentally measured boundary shear and normal forces, full-field displacements, and full-field thickness [85,91,101]. PO and LRT samples generally exhibited reduced stiffness (Figure 5A, B), with the lowest values localized to the infarct core and modest increases found in the adjacent borders. Means were calculated from these full-field distributions and compared using a non-parametric Kruskal-Wallis test and *post-hoc* Dunn’s tests (Figure 5C). PO and LRT samples exhibited a reduction in mean stiffness (sham: 235.0 ± 113 kPa; PO: 137.7 ± 112 kPa) relative to sham samples, although this difference was only significant for the LRT group (100.2 ± 98.7 kPa; *p* < 0.05). Both infarcted groups exhibited modestly increased heterogeneity in stiffness compared to the sham group (with CoV of 0.380 ± 0.12 (sham), 0.566 ± 0.16 (PO), and 0.488 ± 0.13 (LRT)), though only the PO group approached statistical significance relative to the sham group (*p* < 0.10; Figure 5D).

**Figure 5.**
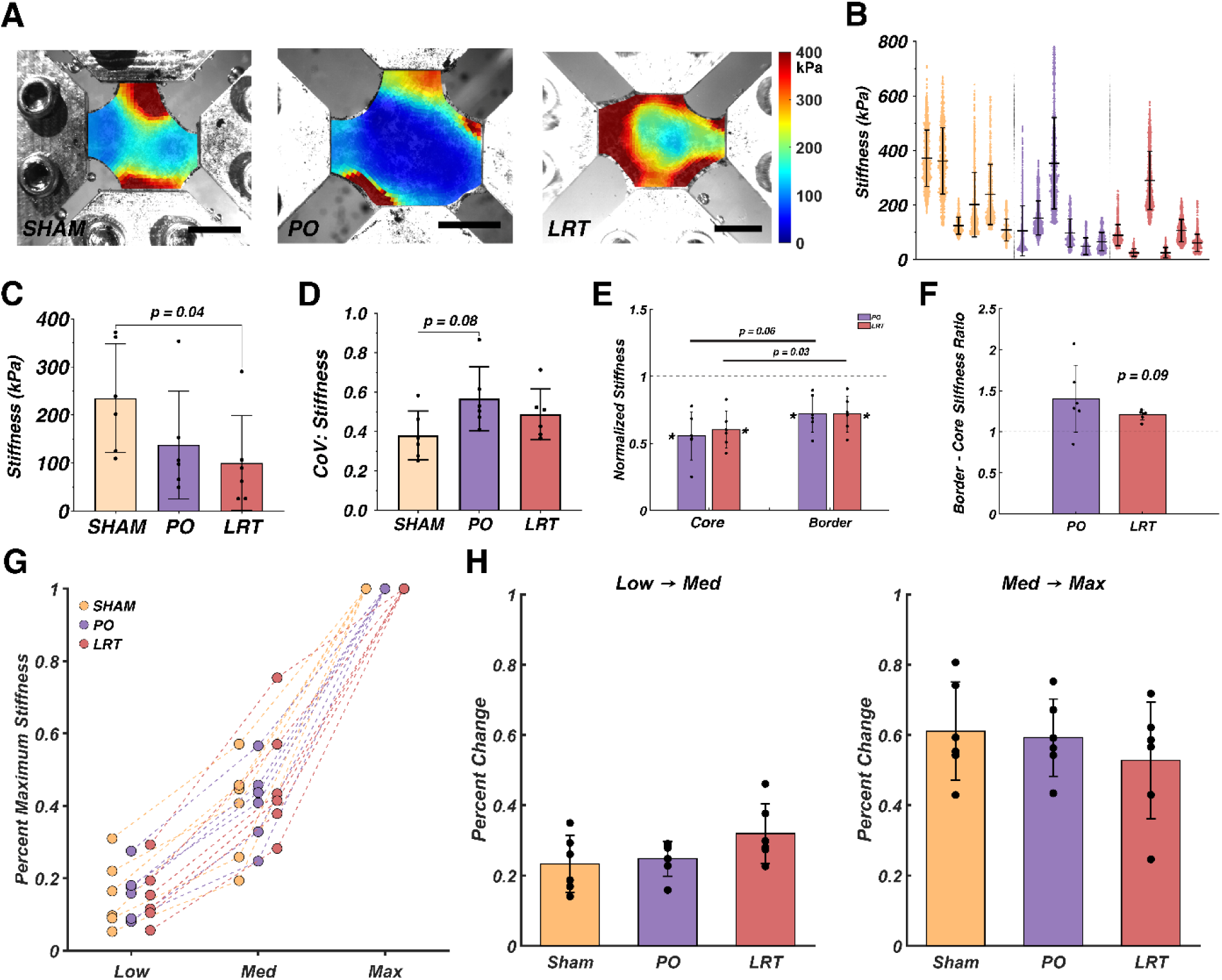
Full-field stiffness distributions (A) for representative sham (left), PO (center), and LRT (right) samples. Scatter plots showing full-field stiffness distributions for all samples tested (B). A bar plot comparing mean stiffnesses (C) and the coefficients of variation (CoV) in stiffness between groups (D). Inter-group statistical significance was detected via a Kruskal-Wallis test with *post-hoc* Dunn’s tests. Infarct core and border stiffnesses normalized to the remote region (E) and the ratio of infarct border-to-core stiffnesses (F) for PO and LRT samples. Intra-group statistical significance was detected via Wilcoxon matched-pairs signed rank test. (G) Stiffness at low (∼7%) and medium (∼14%) boundary strain levels for sham, PO, and LRT samples normalized to stiffness at peak strain (∼22%). (H) The percent change in normalized stiffness for the low-medium strain transition and the medium-maximum strain transition. No inter-group statistical significance was detected via a Kruskal-Wallis test. An asterisk (*) denotes *p* < 0.05 when regional values were compared to a remote value of one using a Wilcoxon signed-rank test. An octothorpe (#) denotes *p* < 0.10 for a Wilcoxon signed-rank test. Scale bars indicate 5 mm.

When infarct core and border stiffnesses were normalized to each sample’s remote mean value, a Wilcoxon signed-rank test showed that the normalized infarct core stiffness was significantly lower than a value of one for both PO and LRT groups (PO: 0.556 ± 0.18, *p* < 0.05; LRT: 0.601 ± 0.14, *p* < 0.05; Figure 5E). This trend was maintained when we compared normalized border stiffness to a value of one (PO: 0.723 ± 0.14, *p* < 0.10; LRT: 0.720 ± 0.13, *p* < 0.05; Figure 5E). A Wilcoxon matched-pairs signed rank test revealed that the borderzones of both PO and LRT samples were stiffer than their respective infarct cores, but this difference was only significant for the LRT group. The border-to-core stiffness ratio, indicating the transition in mechanical properties between these adjacent regions trended lower in the LRT group (Figure 5F; PO: 1.40 ± 0.41; LRT: 1.21 ± 0.06; *p* < 0.10; Figure 5F). Additionally, it was markedly more consistent in LRT samples (CoV = 0.05) compared to PO samples (CoV = 0.29). PO samples spanned extremes from a border that was 15% less stiff than the infarct core to a border nearly 200% stiffer than the infarct core. In contrast, LRT samples had borderzones that were consistently 10 – 20% stiffer than their infarct cores.

To examine the effects of MI on nonlinear stress-strain behavior, we implemented orthotropic GAIM by inputting forces and displacements from earlier in each extension (boundary strains of ∼7% and ∼14%). After normalizing each sample’s mean stiffness to its stiffness at peak load (boundary strain of 22%), we used a Kruskal-Wallis test to detect statistical differences in the percent change in normalized stiffness for the low-medium strain transition and the medium-maximum strain transition between groups (Figure 5G, H). All groups exhibited similar increases in stiffness during the low-medium transition (sham: 23 ± 8.1%; PO: 25 ± 5.0%; LRT: 32 ± 8.5%) and the medium-maximum transition (sham: 61 ± 14%; PO: 59 ± 11%; LRT: 53 ± 17%), and there were no statistical differences detected between groups. This prompted us to conclude that the sham, PO, and LRT groups all exhibited similar levels of nonlinearity during biaxial testing.

### 3.4 LRT Does Not Alter Local Ex Vivo Strain Distributions

We utilized DIC [84,102] to calculate full-field Green-Lagrange strain tensors for sham, PO, and LRT samples during planar biaxial testing. Figure 6A shows the full-field Green-Lagrange strain for representative sham, PO, and LRT samples at peak equibiaxial extension and Figure 6B shows the full-field distributions for each sample tested. Mean horizontal (*Exx)* and vertical (*Eyy*) strains were comparable between sham, PO, and LRT groups (*Exx*: sham: 0.134 ± 0.009; PO: 0.111 ± 0.05; LRT: 0.127 ± 0.02; Figure 6C, left; *Eyy*: sham: 0.149 ± 0.04; PO: 0.131 ± 0.03; LRT: 0.156 ± 0.0; Figure 6E, left). Compared to the sham group, the infarcted groups (PO and LRT) demonstrated modestly increased variability in both normal strain directions (CoV of *Exx*: sham: 0.509 ± 0.24; PO: 0.988 ± 1.1; LRT: 0.898 ± 0.27; Figure 6C, right; CoV *of Eyy*: sham: 0.281 ± 0.13; PO: 0.383 ± 0.16; LRT: 0.431 ± 0.27; Figure 6E, right). However, these increases did not reach statistical significance. Across groups, mean shear strains (*Exy*) were consistently low (near zero, Figure 6B) and spatially variable during this extension, resulting in unreasonably large CoVs, and were omitted from the analysis presented here, but are included in the supplement (Figures S9 – S11).

**Figure 6.**
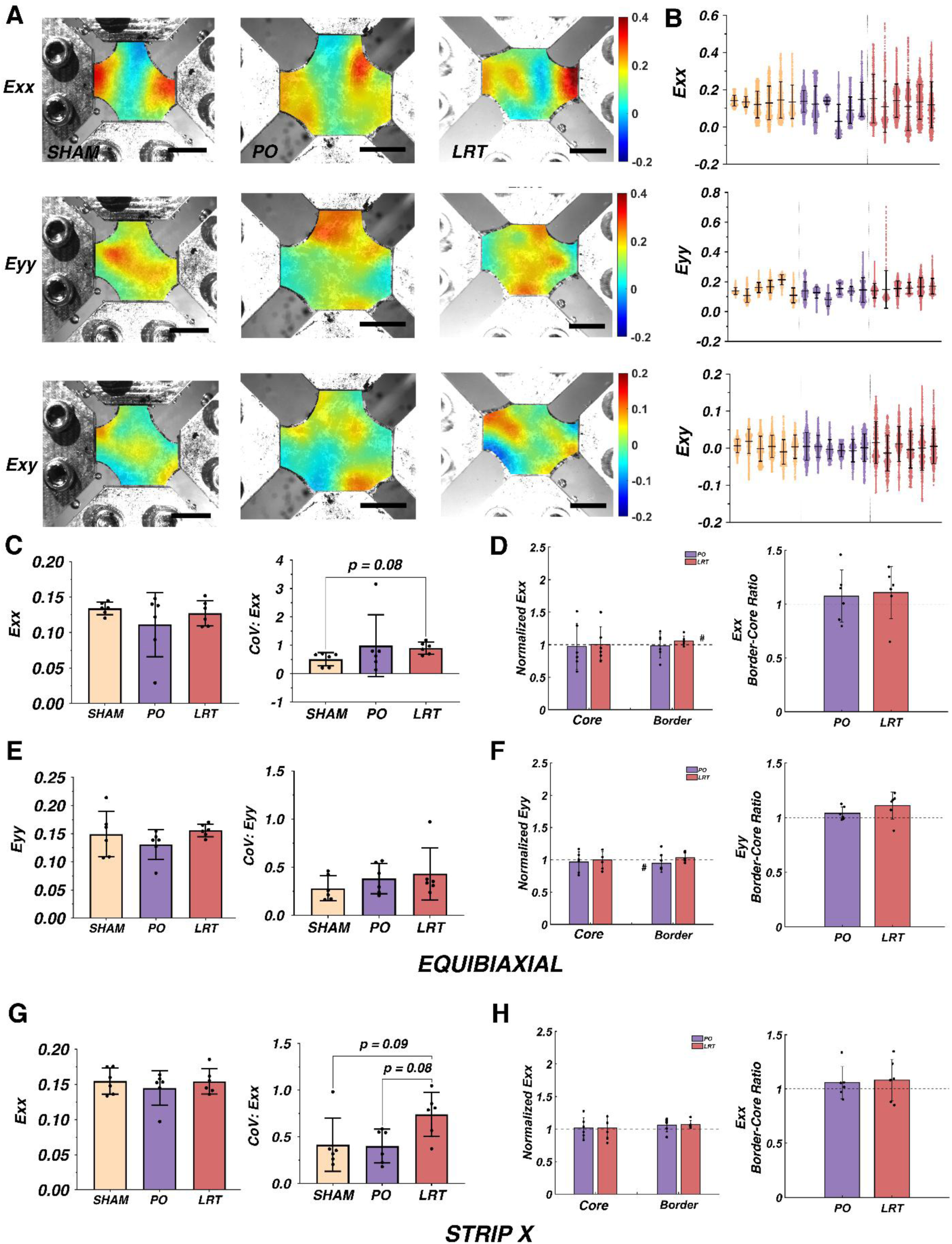
Green-Lagrange strain fields (A) at peak equibiaxial extension for a representative sham (left), PO (center), and LRT (right) sample. (B) Scatter plots showing full-field strain distributions at peak equibiaxial extension all samples tested. (C) Bar plots comparing mean horizontal strain (*Exx*; left) and the coefficients of variation (CoV) in *Exx* (right) at peak equibiaxial extension between the sham, PO, and LRT groups. (D) Normalized infarct core and border *Exx* (left) and border-to-core *Exx* ratios (right) at peak equibiaxial extension. (E) Bar plots comparing mean vertical strain (*Eyy*; left) and the CoV in *Eyy* (right) at peak equibiaxial extension. (F) Normalized infarct core and border *Eyy* (left) and border-to-core *Eyy* ratios (right) at peak equibiaxial extension. (G) Bar plots comparing mean *Exx* (left) and the CoV in *Exx* (right) at peak Strip X extension. (H) Normalized infarct core and border *Exx* (left) and border-to-core *Exx* ratios (right) at peak Strip X extension. Inter-group statistical significance was detected via a Mann-Whitney *U*-test. An asterisk (*) denotes *p* < 0.05 when regional values were compared to a remote value of one using a Wilcoxon signed-rank test. An octothorpe (#) denotes *p* < 0.10 for a Wilcoxon signed-rank test. Scale bars indicate 5 mm.

There were no prominent differences in normalized regional strain distributions between groups (PO vs. LRT) or regions (infarct core vs. border) during the equibiaxial extension (Figure 6D, F). Additionally, Wilcoxon signed-rank tests revealed few differences between normalized infarct core and border values and the remote region. In the LRT group, borderzone *Exx* values trended toward a significant increase relative to the normalized remote value of one (*p* < 0.10). Meanwhile, in the PO group, borderzone *Eyy* values trended towards a decrease (*p* < 0.10). To further explore the effects of infarction and LRT on regional Green-Lagrange strain fields, we repeated the analysis for *Exx* values during the Strip X extension, which primarily loads circumferentially-aligned fibers (Figure 6G, H; [108,109]). Normalized horizontal strain (*Exx*) was similar between PO and LRT samples (Figure 6H, right), with no evidence of regionally-dependent differences (Figure 6H, left). Normalized vertical strains (*Eyy*) were small during the Strip X extension, complicating the calculation of CoV and normalized values, and are presented in the supplement (Figure S12).

### 3.5 Structural Anisotropy Strength is Higher in the Border than the Infarct Core

Dynamic QPLI [86] captured full-field DoLP, a measure of the strength of collagen fiber alignment, throughout planar biaxial testing. Full-field DoLP distributions were similar in sham, PO, and LRT samples in the unloaded state and at peak equibiaxial extension (Figure 7A, B). There were no significant differences in the loaded mean DoLP distributions (sham: 0.228 ± 0.04; PO: 0.237 ± 0.03; LRT: 0.253 ± 0.02) or the CoVs in loaded DoLP (sham: 0.464 ± 0.06; PO: 0.436 ± 0.07; LRT: 0.441 ± 0.03) between groups during the equibiaxial extension (Figure 7C). CoVs were high, however, across all experimental groups, indicating prominent spatial heterogeneity. The change in DoLP (ΔDoLP; loaded – unloaded DoLP) at peak equibiaxial extension was higher in the infarcted samples than sham samples (sham: 0.0270 ± 0.009; LRT: 0.0526 ± 0.008; PO: 0.0420 ± 0.04), but this difference was only significant for LRT samples (Figure 7D, left; *p* < 0.05). Notably, ΔDoLP was also less spatially variable in the LRT group than in the sham group (Figure 7D, right; sham CoV: 2.21 ± 0.47; PO CoV: -0.276 ± 4.83; LRT CoV: 1.45 ± 0.21; *p* < 0.05 between sham and LRT).

**Figure 7.**
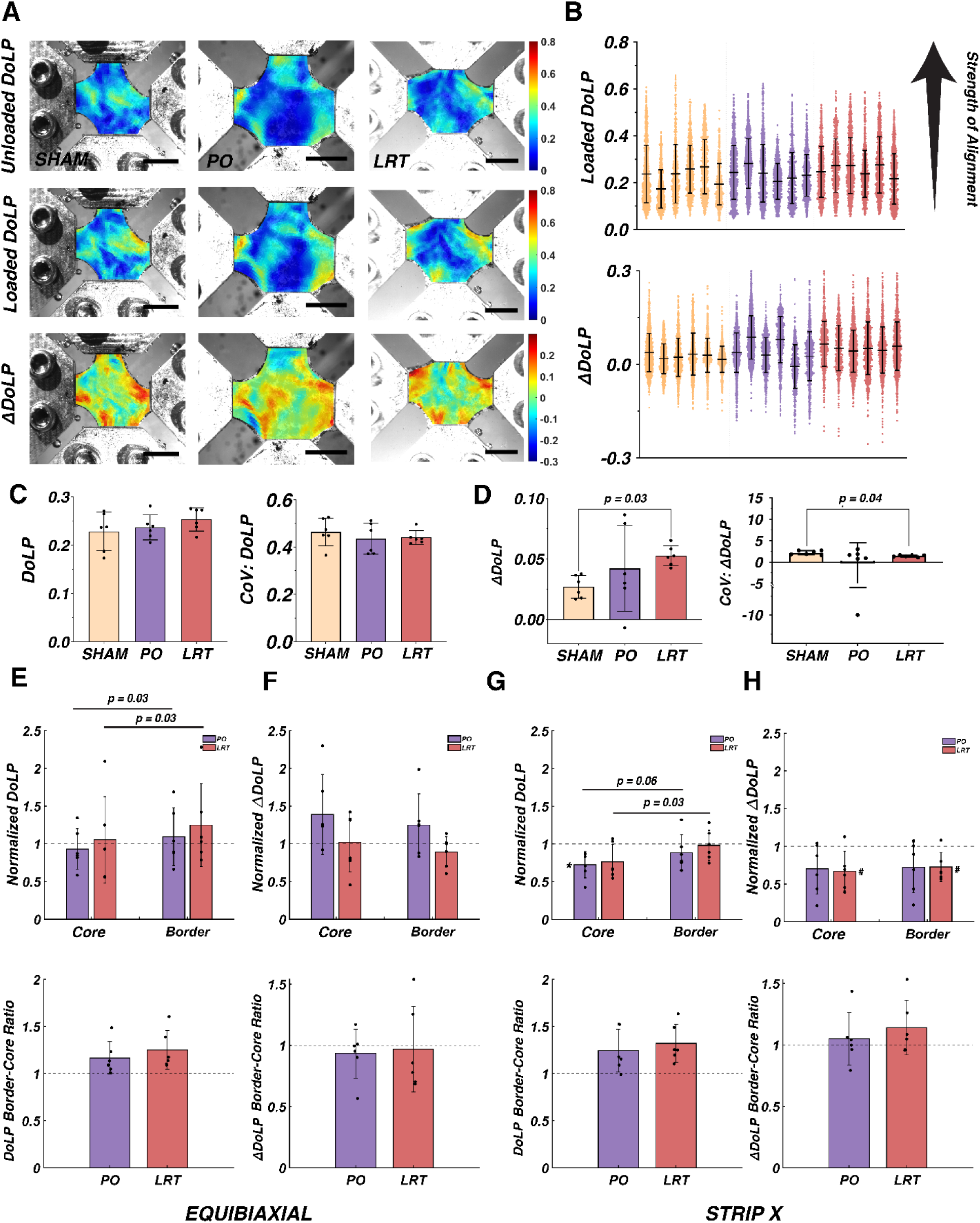
QPLI DoLP fields (A; unloaded, loaded, and ΔDoLP) at peak equibiaxial extension for representative sham (left), PO (center), and LRT (right) samples. (B) Scatter plots showing full-field DoLP distributions at peak equibiaxial loading (B, top) and the distribution of the ΔDoLP during equibiaxial loading for all samples (B, bottom). (C) Bar plots comparing mean DoLP values (left) and the coefficient of variation (CoV) in DoLP (right) at peak equibiaxial loading between sham, PO, and LRT groups. (D) Bar plots comparing mean ΔDoLP values (left) and the CoV in ΔDoLP (right) at peak equibiaxial loading between sham, PO, and LRT groups. Inter-group statistical significance was detected via a Kruskal-Wallis test with *post-hoc* Dunn’s tests. (E) Normalized infarct core and border DoLP values (top) and the border-to-core DoLP ratios (bottom) at peak equibiaxial extension. (F) Normalized infarct core and border ΔDoLP (top) and the border-to-core ΔDoLP ratios (bottom) at peak equibiaxial extension. (G) Normalized infarct core and border DoLP values (top) and border-to-core DoLP ratios (bottom) at peak Strip X extension. (H) Normalized infarct core and border ΔDoLP (top) and border-to-core ΔDoLP ratios (bottom) a peak Strip X extension. For panels E-H, intra-group statistical significance is shown for a Wilcoxon matched-pairs signed rank test and inter-group statistical significance is shown for a Mann-Whitney U-test. An asterisk (*) denotes *p* < 0.05 when regional values were compared to a remote value of one using a Wilcoxon signed-rank test. An octothorpe (#) denotes *p* < 0.10 for a Wilcoxon signed-rank test. Scale bars indicate 5 mm.

Normalized DoLP at peak equibiaxial extension in infarct cores was significantly smaller than that of infarct borders for both PO and LRT samples (Figure 7E, top; PO: 0.93 ± 0.27; LRT: 1.05 ± 0.57). Thus, the border-to-core DoLP ratio was greater than unity, but did not differ between groups (Figure 7E, bottom; PO: 1.16 ± 0.17; LRT: 1.25 ± 0.21). Next, we compared normalized ΔDoLP values at peak equibiaxial extension (Figure 7F, top). While PO samples (infarct core: 1.39 ± 0.53; border: 1.25 ± 0.41) exhibited modestly larger normalized ΔDoLP values than LRT samples (infarct core: 1.02 ± 0.39; border: 0.893 ± 0.20) in both infarct cores and borders, this was not statistically significant. The border-to-core ΔDoLP ratio also did not differ between groups (Figure 7F, bottom; PO: 0.933 ± 0.20; LRT: 0.970 ± 0.35). Neither the PO nor LRT groups exhibited statistically significant differences from the remote value of one for normalized DoLP or ΔDoLP in either the infarct core or borderzone at peak equibiaxial extension.

We repeated this analysis for the Strip X biaxial extension and observed trends similar to the equibiaxial extension. Normalized DoLP at peak Strip X extension was lower in the infarct core compared to the border for both groups, with this difference reaching significance in the LRT group (Figure 7G, top; core: 0.762 ± 0.23, border: 0.980 ± 0.20, *p* < 0.05) and approaching statistical significance in the PO group (Figure 7G, top; core: 0.724 ± 0.17, border: 0.886 ± 0.24, *p* < 0.10). Again, this yielded a border-to-core DoLP ratio greater than unity (Figure 7G, bottom; PO: 1.24 ± 0.23; LRT: 1.32 ± 0.20), which did not differ between the PO and LRT groups. Furthermore, normalized ΔDoLP values were similar between PO and LRT groups and did not differ regionally (Figure 7H, top). Normalized ΔDoLP border-to-core ratios also did not differ between groups (Figure 7H, bottom). Wilcoxon signed-rank tests showed that PO samples had statistically significantly lower normalized DoLP values in the infarct core than the periphery (Figure 7G; *p* < 0.05), whereas LRT samples showed a trend toward decreased ΔDoLP values in both the infarct core and border (Figure 7H; *p* < 0.10 for both regions).

## 4. Discussion

LRT is well-established as one of the most effective strategies for reducing rates of post-MI ventricular rupture [14,26–28], but how it protects the heart, alters spatial heterogeneity, and impacts the dynamic organization of the LV ECM is unknown [3]. To address this gap, we utilized novel, full-field experimental techniques to quantify the geometric [84], mechanical [85], and structural [86] properties of infarcted LV ECM. When compared to PO, LRT generated a larger infarct borderzone (Figure 2). Furthermore, the borderzone area of LRT samples remained larger than that of PO samples under a variety of biaxial extensions. We also found that infarction exacerbated geometric heterogeneity and increased overall sample thickness, mainly due to thickening in the infarct core and, to a lesser extent, in the infarct border (Figure 4). However, no significant differences in thickness were observed between PO and LRT samples. Full-field stiffness results from our GAIM method[85] revealed that infarction reduced stiffness, with the most prominent reductions in the infarct core, followed by the infarct border (Figure 5). Notably, LRT promoted a more gradual and consistent transition in stiffness from the infarct core to the remote ECM (Figure 5E, F). Finally, we showed that infarction does not significantly alter local strain distributions during equibiaxial and Strip X extensions (Figure 6), and that while infarction locally reduced DoLP values in the infarct core vs. the border region, LRT does not further modulate this reduction (Figure 7E, G). Our findings that LRT broadened the border between the infarct core and remote tissue and created a more gradual and uniform transition in geometric and mechanical properties throughout the heart suggest *in vivo* ventricular rupture may be limited by a reduction in stress concentrations within the infarct core and border, two of the most common sites of ventricular rupture in animal models [14,65,69,82,83]. This work represents a step towards a more mechanistic understanding of how LRT protects the infarcted heart and highlights the importance of full-field experimental techniques for clarifying how MI alters the mechanical and structural properties of the LV free wall ECM. Together, these contributions refine our overall understanding of post-MI healing, define how LRT alters this process, and may guide the development of next-generation post-MI therapies.

### 4.1 LRT Creates a Larger Borderzone

In post-MI literature, the definition of infarct borderzones remains controversial [3,21,72–77]. As far back as the 1960s, there are reports of moderately damaged myocardium surrounding the necrotic infarct core during the first week post-MI [72,73,110]. More recently, Berry et al. [75] utilized atomic force microscopy to estimate the spatial distribution of the elastic modulus of infarcted myocardium. Although these measurements were obtained at two weeks post-MI, long after inflammation and rupture risk have subsided, they depicted a clear transitional region (∼5 mm in width) from the stiff infarct core to the more compliant remote myocardium. Similarly, spatial transcriptome analyses of murine infarcts revealed the presence of a narrow borderzone surrounding infarct cores within the first week post-MI [111]. These borderzones were characterized by an upregulation of mechanosensing genes that may regulate LV remodeling [76]. These more modern studies, which utilized advanced imaging and mechanical testing modalities, have provided evidence in support of a biologically and mechanically distinct infarct borderzone. However, these studies utilized PO models of MI, leaving the effects of RT on the size and properties of the borderzone undefined. As RT and LRT are more clinically relevant models of MI [24,54–57], a more mechanistic understanding of the relationship between RT timing, borderzone size and properties, and ventricular rupture risk offers a path forward for improving therapeutic interventions and MI patient outcomes.

Like many early studies [24,54–57], we found that LRT did not reduce infarct size. There were no differences in the area of the infarct core (Figure 2C), the ratio of the infarcted tissue area (infarct core + border) relative to total sample area (Figure 2E), or the ratio of the infarct core area relative to the total sample area (Figure S7F) between the PO and LRT groups. In contrast, LRT promoted a larger borderzone (Figure 2D) and greater border-to-core area ratios (Figure 2F) when compared to PO. Given past reports of increased rates of rupture in the borderzone [14,26–28], our observations of a larger border region may initially suggest LRT should exacerbate risk. That said, failure mechanics of simpler materials show that tears initiate and propagate in regions of intense stress concentration. The infarct borderzone is a site of prominent heterogeneity and we observed that the border broadening associated with LRT reduced spatial transitions in thickness, potentially decreasing stress concentrations in vulnerable regions of the LV (Figure 4).

### 4.2 LRT Promotes a Gradual Increase in Stiffness Between Infarct Core and Remote ECM

Overall, we observed a decrease in the stiffness of the LV ECM with infarction (Figure 5C). This observation aligns with the passive uniaxial testing results from Connelly et al. [69], particularly for the LRT group. In their study, which utilized a rabbit model of MI, LRT samples were approximately 30% less stiff (defined in their study as tensile force/thickness) than PO samples during the first two days of post-MI healing [69]. In the present study, mean spatial stiffness was 31.5% lower for the LRT group relative to the PO group. Unlike their study, however, we also observed lower mean stiffnesses (-52.2%) in the PO group than the sham group, rather than the higher (+25%) stiffnesses reported by Connelly et al. [69] at this time point. These differences could arise from discrepancies between different animal models of MI and rates of post-MI healing, differing methods of thickness measurement [84,112,113], or their use of uniaxial testing on homogeneous longitudinally-oriented strips of cellularized infarct core tissue.

One of the most novel contributions of this study is our use of a full-field inverse technique, which was previously validated for a variety of simulated and experimental datasets [85,101,114]. This approach, alongside the common *k*-means clustering algorithm [105,106], allowed us to quantify regional changes in stiffness within the infarct core, infarct border, and remote LV ECM. In general, stiffness was lowest in the infarct core, followed by the border, and then the remote tissue (Figure 5E). We found that LRT decreased the transition in stiffness across the infarct border and that the borders of LRT samples were 1.21 ± 0.06 times as stiff as their cores, whereas PO samples had borders that were 1.40 ± 0.41 times as stiff (Figure 5F). This transition was also markedly more consistent – LRT sample infarct borders were 10 – 20% stiffer than their infarct cores, whereas PO sample borders were anywhere from 15% less stiff to 200% stiffer than their respective infarct cores. Together, these results suggest that LRT promotes a more gradual and regular transition in stiffness between the infarct core and the peripheral remote LV ECM.

Another innovative aspect of this study afforded by full-field measurements was quantification of heterogeneity. Sham samples were relatively homogeneous in thickness (Figure 4D) and stiffness (Figure 5D), but Green-Lagrange strain fields were highly heterogeneous in sham, PO, and LRT samples (Figure 6C, E, G). Even for symmetric extensions, such as the equibiaxial extension, mean CoV in strain at peak extension did not differ between sham, PO, and LRT groups and there were no regional differences in strain between or within PO and LRT groups (Figure 6D, F, and H). In PO and LRT samples, spatial variability was increased for both thickness and stiffness (Figures 4DG and 5DE). Increased heterogeneity in LV ECM geometry and mechanics could exacerbate the stress concentrations in the heart wall as it is being loaded *in vivo*. Accentuated stress concentrations were something that Connelly et al. [69] noted during *ex vivo* failure testing of intact PO and LRT hearts. Their estimates of wall stress highlighted increased stresses in the borders of ruptured PO hearts, suggesting a link between rupture likelihood and local stress. In contrast, the larger borders (Figure 2D, F, G-J) and subtler transitions in geometry (Figure 4G) and mechanical properties (Figure 5F) following LRT, as reported here, may serve a protective role by broadening and mechanically smoothing the borderzone during post-infarction inflammation. Both *ex vivo* and *in vivo* estimations of full-field stress and failure testing are needed to confirm the theory that these alterations translate to a reduction in stress concentration and an increase in overall failure strength.

### 4.3 LRT Does Not Alter Collagen Density or Alignment

DoLP distributions, which indicate local strength of collagen fiber alignment, were highly heterogeneous across the three groups (Figure 7). Regional comparisons of normalized DoLP indicated lower collagen fiber alignment in infarct cores compared to borders (Figure 7E, G). However, there were no differences between PO and LRT samples in infarct core or border normalized DoLP or the normalized ΔDoLP (Figure 7E – H). Thus, our use of full-field, dynamic QPLI did not detect differences in collagen fiber anisotropy strength between PO and LRT groups. This suggests that LRT, as compared to PO, has a minimal effect on the overall disorganization of collagen fibers in the LV at this time point in post-MI healing. These results, in addition to picrosirius red staining (Figure S3), are consistent with several past studies that found no discernible differences in bulk collagen content between homogeneous samples taken from the infarct cores of PO and LRT samples [62–64].

To further contextualize the spatial variability in the organization of collagen fibers measured via QPLI, we used non-polarization-resolved second harmonic generation (SHG) imaging to visualize collagen fibers within a sham, PO, and LRT sample (Figure 8). SHG microscopy is widely used across various tissue types to image collagen fibers (primarily types I and III in the heart; [115]). It is particularly useful for quantifying structural changes in collagen fibrils and fibers associated with pathological conditions, such as fibrosis and connective tissue disorders, due to its nonlinear optical sensitivity to collagen fiber organization. To prepare samples for SHG, we decellularized one sham, PO, and LRT heart as described in the Methods section and shown in Figure 1A, B. Next, LVs were separated from the rest of the heart (Figure 1 C, D) and trimmed to a circumferentially-oriented rectangular strip that included both infarct and remote LV ECM for the PO and LRT groups, and healthy free wall LV ECM for the sham group. Each strip was embedded in an agar gel (2.5 – 4% agarose w/v) and then cut into ∼200 micron-thick slices using a vibratome (Leica VT 1200S; Leica Biosystems, IL). Each slice was then laid flat on a glass slide, hydrated with PBS, covered with a coverslip, and sealed using nail polish. The prepared sample was then placed under a microscope (20x objective, zoom2; Olympus Fluoview 300 laser scanning system mounted on an Olympus BX61 upright microscope stand; see Chen et al. for a comprehensive overview [116]) and imaged via non-polarization-resolved SHG with circularly polarized light, which excites collagen fibers equally, regardless of their orientation. We used an excitation wavelength of 780 nm and collected the SHG signal by placing a 390/18 filter before both forward and backward photomultiplier tubes. After collecting each image, we then projected the forward and backward channel of each SHG image and analyzed the direction of collagen fiber alignment, fiber width, and fiber length using CT-Fire and default image analysis settings [117]. We also quantified porosity as the ratio of void space to total volume in a material using a technique originally implemented to analyze porous tissue engineering scaffolds and biomaterials [118,119].

**Figure 8.**
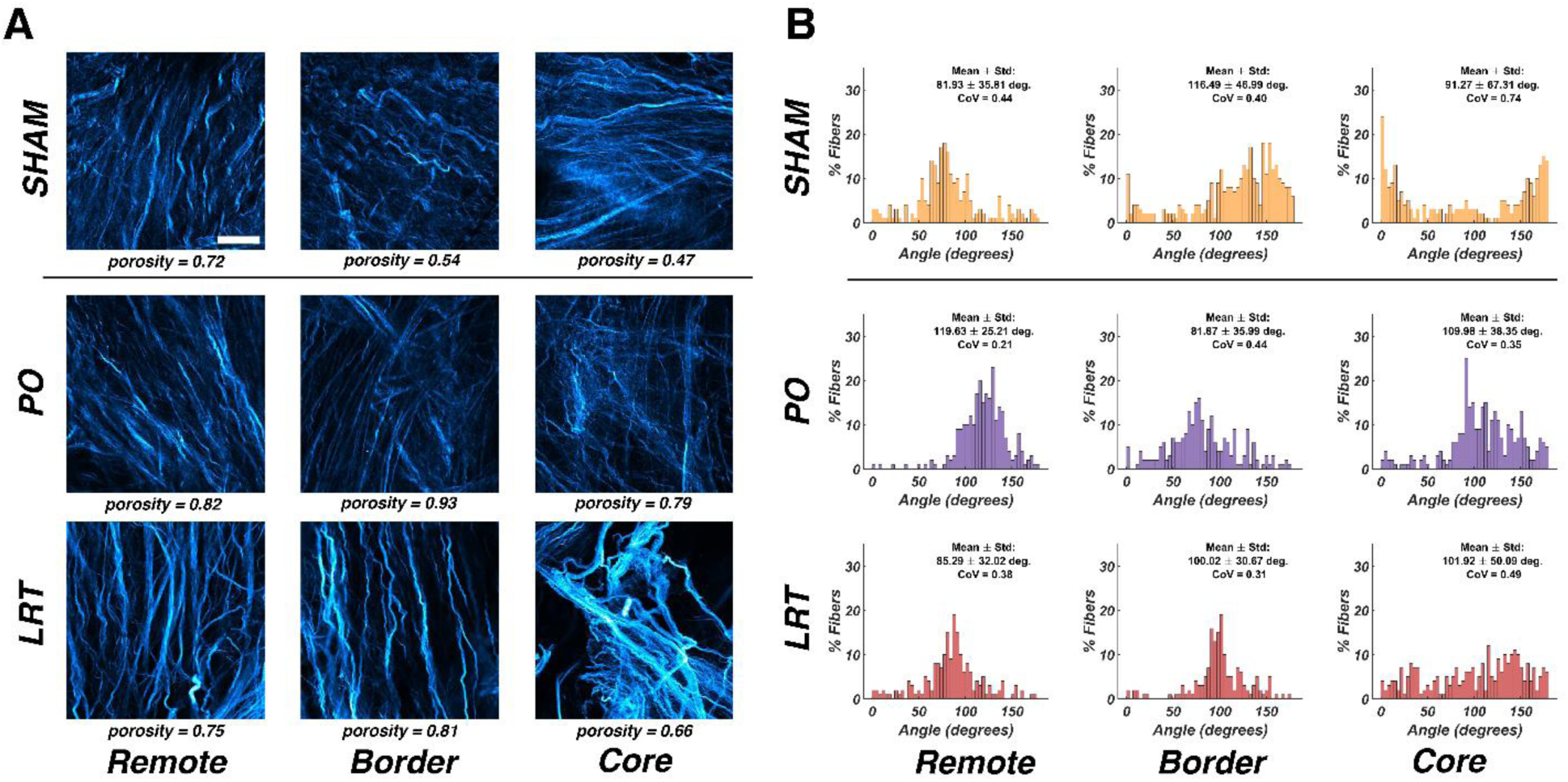
(A) Second harmonic generation (SHG) images of collagen fibers in decellularized sham, PO, and LRT samples. Images for PO and LRT samples are from the remote, infarct border, and infarct core regions. (B) Corresponding collagen fiber direction distributions from CT-Fire showing the mean, SD, and COV. All SHG images are 350 x 350 μm in size. Scale bar indicates 70 μm.

SHG imaging revealed subtle differences in the organization of collagen fibers between treatment groups, particularly at the infarct core-border interface (Figure 8A). In each image from the sham sample, there was a predominant direction of collagen fiber alignment (Figure 8 A, B top row), corroborating observations from Streeter et al. [108] and Pope et al. [109], though the direction of collagen fiber alignment varied with location. The remote regions of the infarcted samples (Figure 8 A, B, left of bottom two rows) appeared similar to the various locations throughout the sham heart in the sense that fibers were relatively organized and had a predominant direction of fiber alignment (PO: 119.6 ± 25° and LRT: 85.3 ± 32° as compared to sham left: 81.9 ± 36°; sham middle: 116.5 ± 47°; sham right: 91.3 ± 67°). In the PO sample, collagen fiber organization was most heterogeneous in the infarct: the CoV was 0.35 and 0.44 in the infarct core and border, respectively, compared to only 0.21 in the remote region. In contrast, the LRT sample demonstrated less heterogeneous collagen fiber alignment in its remote and border regions compared to its infarct core (remote CoV: 0.38; border CoV: 0.31; infarct core CoV: 0.49). Notably, both the LRT and PO samples exhibited a bimodal pattern in porosity with the infarct border having a higher porosity than both the remote and infarct core regions (Figure 8A: PO: 0.93 vs. 0.82 and 0.79; LRT: 0.81 vs. 0.75 and 0.66, respectively). Furthermore, the infarct core of the PO sample exhibited increased porosity compared to the infarct core of the LRT sample (PO: 0.79; LRT: 0.66). These microscale observations mirror the more gradual stiffness transitions we observed in the LRT group (Figure 5F), potentially supporting the notion that LRT preserves local ECM by promoting a more consistent and regular collagenous architecture throughout the infarcted LV free wall. Although these observations are limited by our use of one sample in each group, they offer insight into the heterogenous organization of the ECM in the post-MI heart and how LRT may alter that organization (see Figures S13 and S14 for additional SHG images, as well as Figure S15 for fiber length and width measurements). Also, we would like to note that each SHG image is 350x350 microns in area and captures <1% of the LV ECM free wall, meaning over 1,600 images would be required to visualize the entire surface area of the tissue sample.

### 4.4 Limitations

One major limitation of this study is the small sample size. A total of six hearts per group (sham, PO, and LRT) underwent whole-organ decellularization, dissection, laser micrometry, mechanical testing, and QPLI successfully, and one additional heart per group underwent whole-organ decellularization, dissection, and complementary SHG imaging. It is possible additional samples would address the considerable inter-sample variability observed in many of our measurements. Here, we addressed this variability by normalizing infarct and border means to each sample’s remote mean, enabling us to explore regional differences within each sample. Additionally, we relied on non-parametric statistical tests [120], taking a more conservative approach in our comparisons of groups. Whole-organ decellularization was necessary to isolate the LV ECM for planar biaxial testing, but had the potential to damage the tissue. Ott et al. [90] observed the preservation of both collagen I and III fibers, as well as fiber shape, in decellularized rat hearts using this protocol. They also reported no changes in fiber orientation in decellularized aortic wall and aortic valve leaflets [90]. Additionally, our prior study showed this protocol had minimal overall impact on the mechanical properties of rat right ventricles, provided that thickness differences were properly accounted for in the analyses [91].

Another limitation of our study is the age of the rats used (9 – 12 months), which approximately corresponds to young adulthood in humans (around 25 years of age; [121]). In contrast, MI in humans typically occurs much later in life, with most patients experiencing their first MI between the ages of 65 – 75 [122]. Based on work from Yang et al. [59], which compared ventricular rupture rates in young and aged rodents, we expect that using a more age-appropriate model would exacerbate the mechanical and structural disparities between infarcted and remote tissues, likely due to a more intense immune response. This warrants further exploration, as do sex differences in rupture rates and post-MI outcomes [123,124], especially in the context of therapy design and eventual clinical translation.

Finally, while the experimental techniques applied here were full-field, they did not account for potential variations throughout the different layers of the LV ECM. Planar biaxial testing assumes a state of plane stress and effectively treated the LV ECM as a two-dimensional sample. Full-field Green-Lagrange strain tensors were generated from 2D DIC, which captures only the sample surface deformations. Thus, while GAIM estimations of stiffness incorporated thickness, they quantified heterogeneity *across* the sample surface, but not *throughout* its thickness [84,85]. Similarly, because QPLI relies on transmitted light, it provides summary measures of collagen fiber organization, rather than layer-specific information [86]. Although we expect infarcts to be transmural (extending from the endocardium to the epicardium) based on existing literature [24,46,54], the methods and measurements utilized here are poorly suited to studying variable ECM remodeling throughout the layers of the heart. Nevertheless, we do provide unparalleled spatial information within the circumferential – longitudinal plane of the LV.

### 4.5 Future Directions

This study captures spatial heterogeneity in LV ECM remodeling at one time point in the post-MI healing response (day 1 post-MI), a notoriously temporally dynamic process [3,4,20]. Investigating additional time points later in the inflammatory phase and early in the proliferative phase of post-MI healing (2 – 5 days post-MI) could help clarify how the heart’s geometric, mechanical, and structural properties evolve during healing. Expanding the time course of these experiments would also allow for the evaluation of the potential temporal benefits of LRT (such as accelerated healing) reported by other researchers [56,67–70]. It could also provide new opportunities to identify when and where adverse remodeling is most likely to occur, such that therapeutic interventions have the greatest impact.

Beyond mechanical and structural testing, future studies could also include molecular and cellular-level analyses. For example, immunohistochemical staining to detect shifts in immune cell populations [11,125,126], the fibronectin-dominant provisional matrix [67,127–129], and profiles of matricellular proteins [130,131] and cytokines [132,133] in the various regions of the heart could reveal the extent of inflammation or early fibrotic activity in response to LRT. Single-cell RNA sequencing or transcriptomic profiling may capture transitions in gene expression among important cell types, such as fibroblasts, myofibroblasts, and cardiomyocytes, providing insight into signaling pathways that regulate cell phenotype and subsequent tissue remodeling [111,134,135]. A notable example of this experimental approach comes from Calcagno et al. [111], who used spatial transcriptomics to identify mechanosensitive genes upregulated in borderzone myocardium during the early post-MI period. Likewise, hydrogel-based systems [136] and “infarcts on a chip” have already been used to study how infarct-like oxygen gradients affect cell and tissue behavior [137]. These platforms are valuable tools to study how immune cells and fibroblasts respond to controlled conditions that replicate post-MI healing. Such studies could reveal how cells migrate through the infarcted heart, how the ECM’s heterogeneous structure and mechanics affect them, and how their behavior differs with ischemia and reperfusion. By generating comprehensive datasets on how mechanical and structural properties vary throughout space and time, we can better understand not only which factors most impact rupture risk, but also those that regulate tissue remodeling. This knowledge is key to designing spatially conscious post-MI therapies, such as biomaterials or localized drug delivery systems [138,139], that target the most pertinent regions of the heart at optimal timepoints during post-MI healing. Beyond these technical benefits, these spatiotemporal strategies could improve patient outcomes, particularly for individuals in underserved communities who are less likely to benefit from early RT [49–53].

## 5. Conclusion

LRT substantially improves clinical outcomes following MI, with evidence showing a dramatic reduction in the rates of ventricular rupture [29–36]. Yet, despite this clinical success, the underlying structural and mechanical changes to the LV ECM following LRT remain poorly understood. To investigate how LRT alters post-MI healing, we employed advanced, full-field geometric, mechanical, and structural characterization techniques, including laser micrometry, planar biaxial mechanical testing, and dynamic QPLI. We observed that LRT created a significantly larger infarct borderzone which remained larger in area than PO samples when mechanically loaded (Figures 2 and 3). Infarction also led to marked increases in thickness and geometric heterogeneity (Figure 4). Local tissue stiffness was notably reduced after infarction, and LRT promoted a smoother, more uniform stiffness transition between the infarct core and remote LV ECM, potentially minimizing mechanical discontinuities and stress concentrations (Figure 5). MI did not significantly alter local Green-Lagrange strain distributions in the various regions of the infarcted LV ECM (Figure 6); however, detailed QPLI results indicated reduced collagen fiber alignment within the infarct core in comparison to the border (Figure 7). Complementary SHG imaging supported these observations at the microscale, revealing that infarct cores exhibited increased variability in collagen fiber alignment and that LRT may foster a more gradual transition in collagen fiber organization throughout the infarcted ECM (Figure 8). Collectively, these findings suggest an exciting therapeutic implication: RT, even when administered several hours after an infarction, modifies post-MI remodeling by reducing heterogeneity in the mechanical properties of the borderzone, potentially reducing stress concentrations and ventricular rupture risk. Ultimately, by better defining the full-field, cardioprotective benefits of LRT, our results offer a promising foundation for future studies aiming to develop spatially-tailored therapeutic interventions for improving MI patient outcomes.

## Acknowledgements

This study was funded by a grant from the NSF to CMW (2020173) and from the NIH to PC (1R61CA281795-01). Additionally, we would like to thank Sudhi Chavadam, Michael Chiariello, Elizabeth Gunderson, Mark Nemcek, Shreya Sreedhar, Rhea Nagori, and Simmi Kaur for their help on projects related to this study. The authors would also like to thank the University of Wisconsin School of Medicine and Public Health Biomedical Research Model Services (particularly, Rachel Taylor) for the use of its facilities and their Research Services team for expertise including breeding and specialized microsurgical techniques. Finally, we would like to thank the University of Wisconsin Carbone Cancer Center Experimental Animal Pathology Laboratory (particularly, Toshi Kinoshita) supported by P30 CA014520, for use of its facilities and services and support throughout sample preparation and analysis.

