## Supplement for "Full-Field Analysis Indicates Late Reperfusion Therapy Broadens and Mechanically Smooths the Borderzone During Post-Infarction Inflammation"

**Supplemental Files**

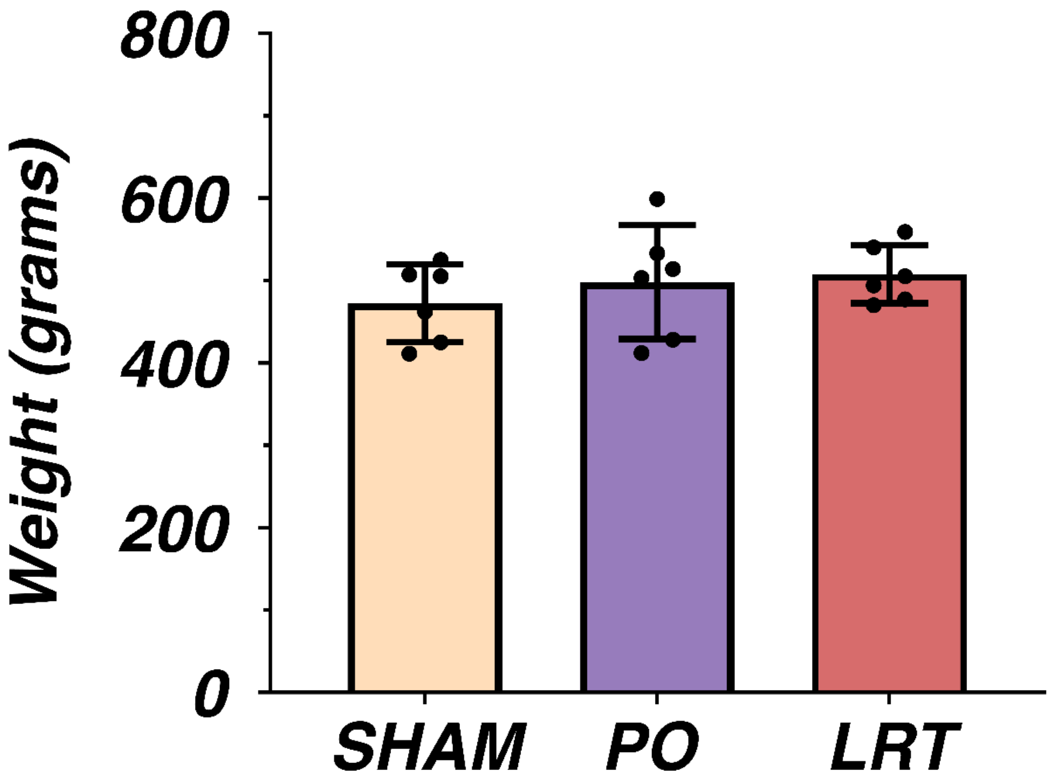


**Figure S1. Rodent pre-surgical body weight (in grams) for the sham, permanent occlusion (PO), and late reperfusion therapy (LRT) groups. No significant differences were detected between groups using a Kruskal-Wallis test (sham: 472.5 ± 47.2 g; PO: 498.2 ± 69.2 g; LRT: 507.5 ± 35.3 g).**

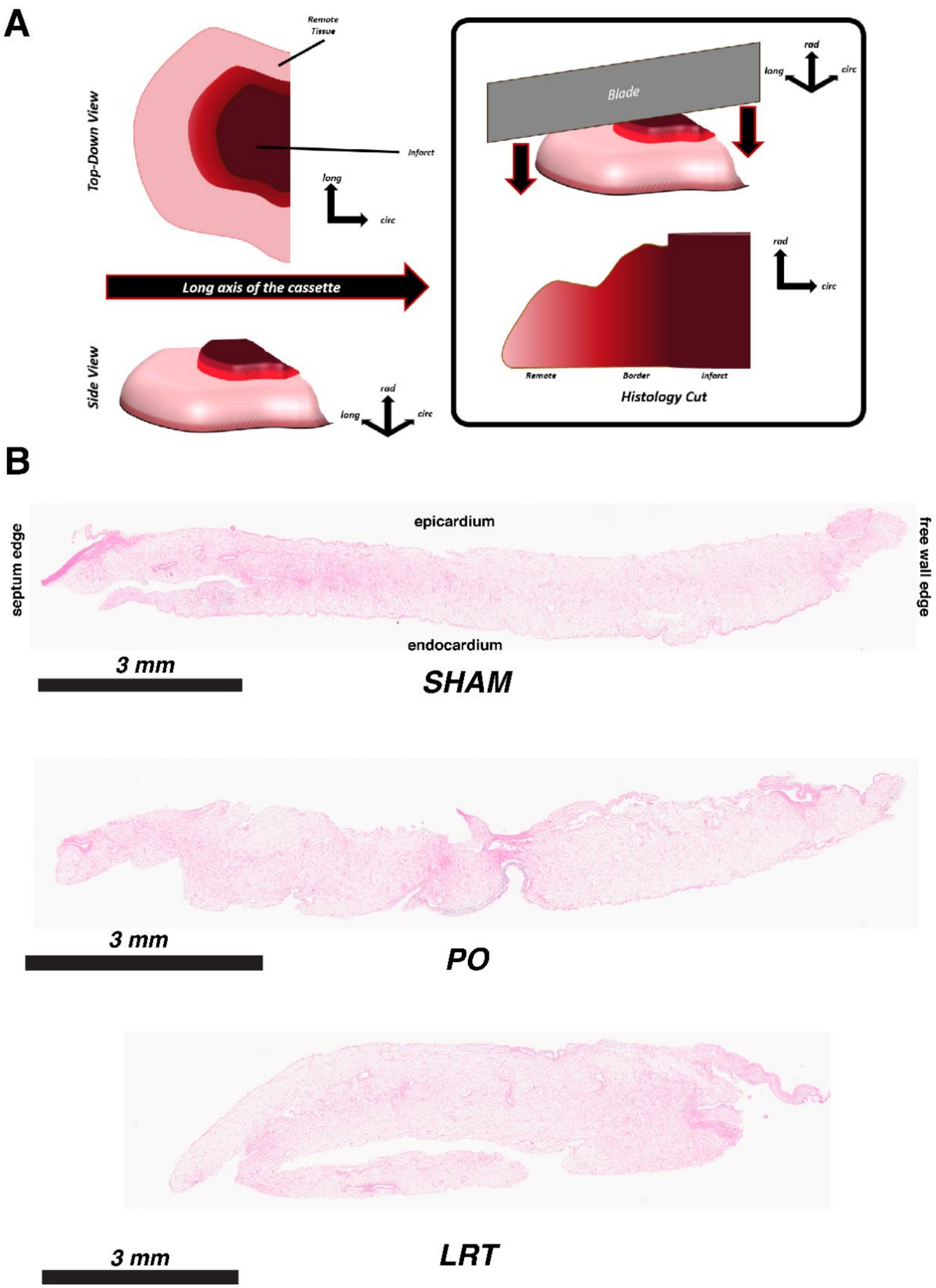


**Figure S2. (A) Schematic showing the orientation of decellularized LV samples and cut plane for histology. (B) Hematoxylin and eosin (H&E) staining performed on 5 micron-thick slices of decellularized rodent left ventricle in representative sham (top), PO (middle), and LRT (bottom) samples. All samples were submerged in formalin for 24 hours, then stored in ethanol. Samples were then cut into 5 micron thick slices, stained, and imaged using a full-slide Aperio scanning microscope at 20x.**

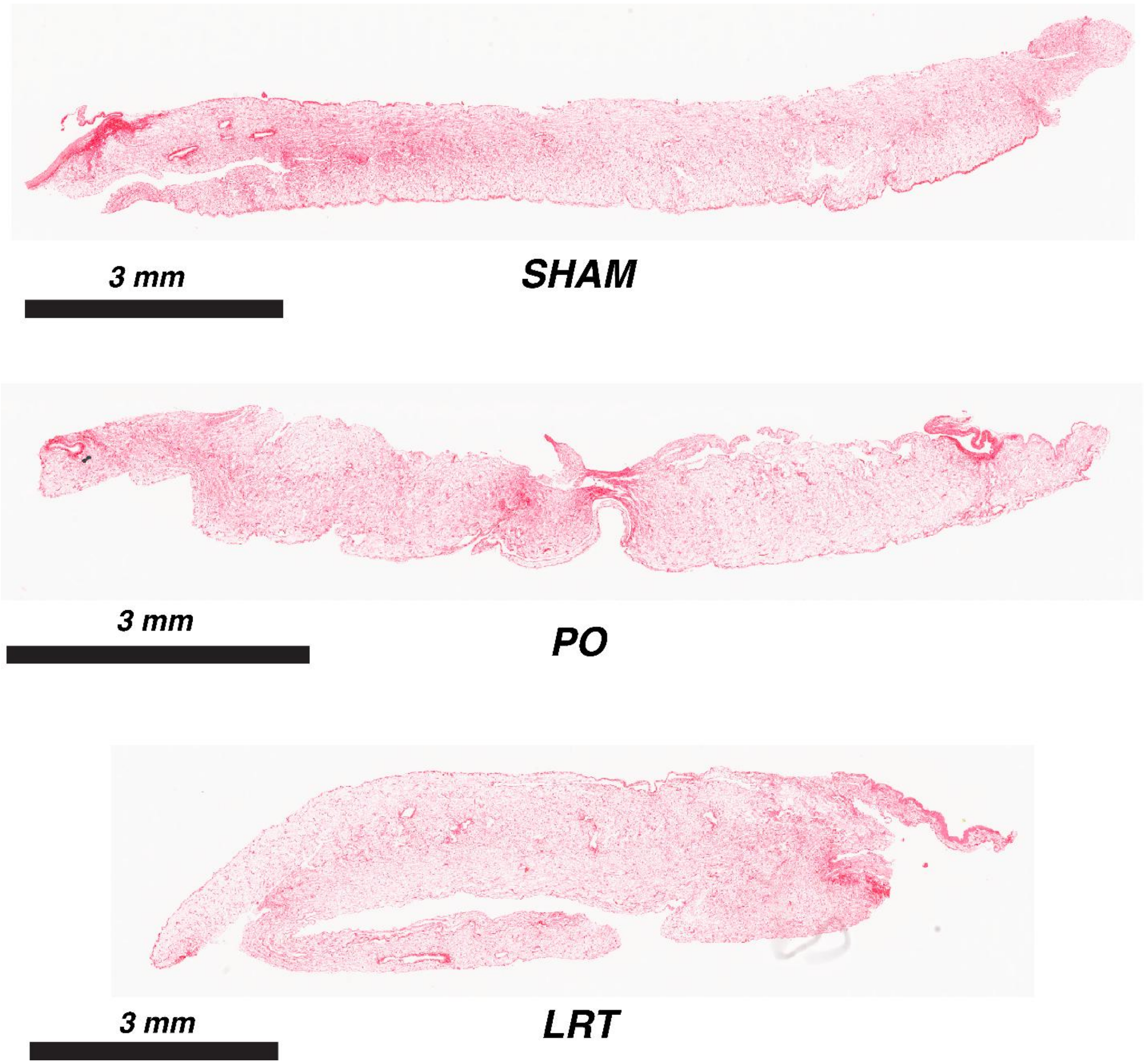


**Figure S3. Picrosirius red (PSR) staining performed on 5 micron-thick slices of decellularized rodent left ventricle in representative sham (top), PO (middle), and LRT (bottom) samples.**

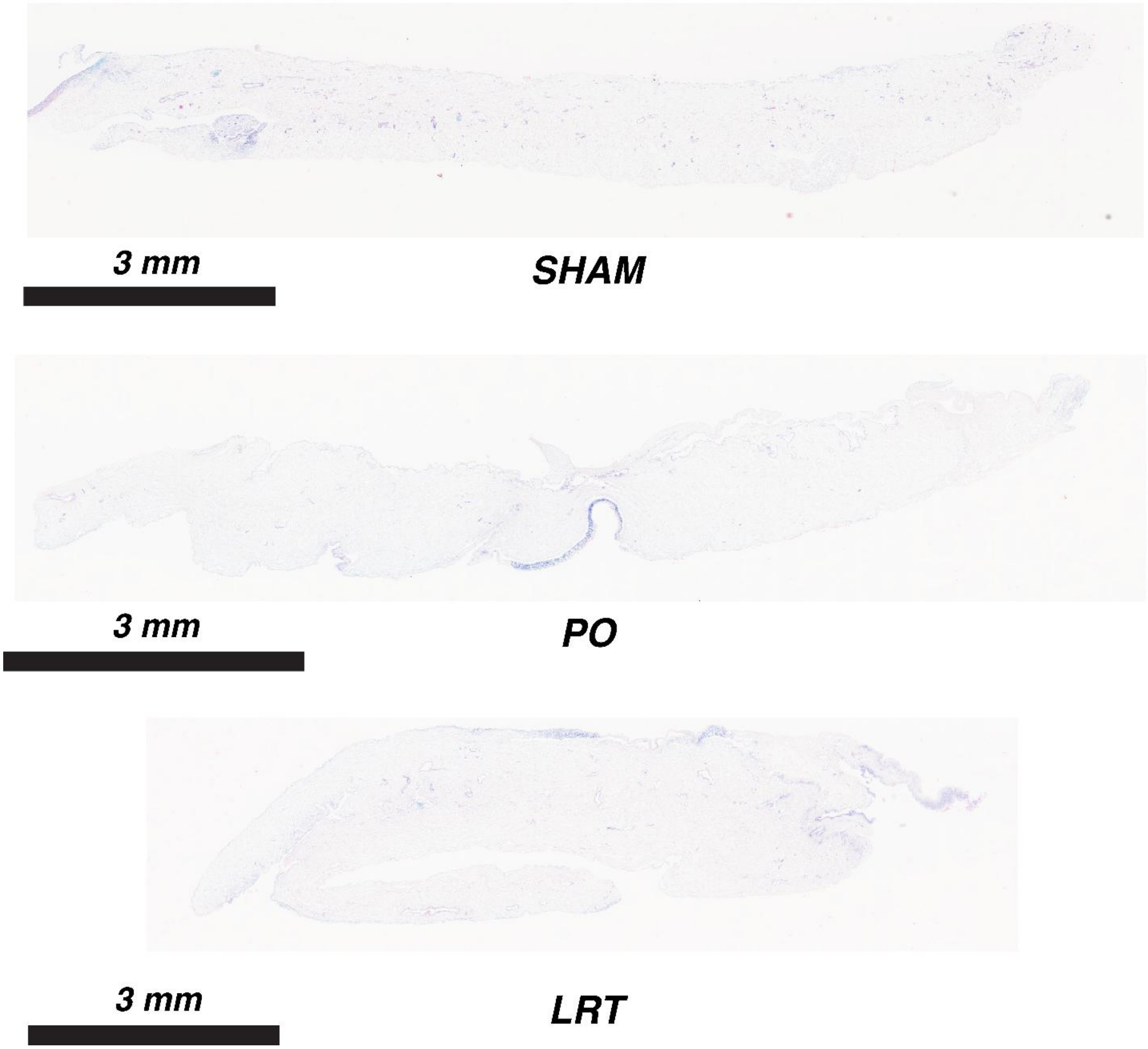


**Figure S4. Alcian blue staining performed on 5 micron-thick slices of decellularized rodent left ventricle in representative sham (top), PO (middle), and LRT (bottom) samples.**

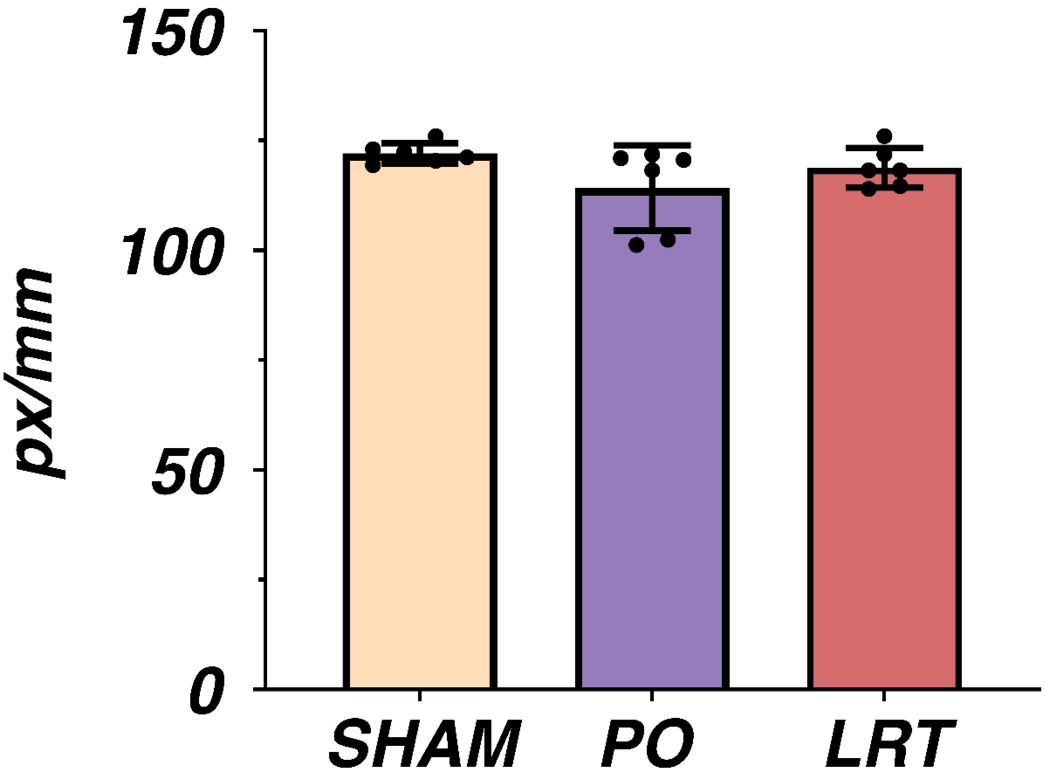


**Figure S5. Quantitative polarized light imaging (QPLI) spatial resolution (in pixels per millimeter) for sham, PO, and LRT groups. Bars represent mean ± SD. No significant differences were detected between groups using a Kruskal-Wallis test (sham: 122.1 ± 2.3 px/mm; PO: 114.2 ± 9.7 px/mm; LRT: 118.8 ± 4.5 px/mm).**

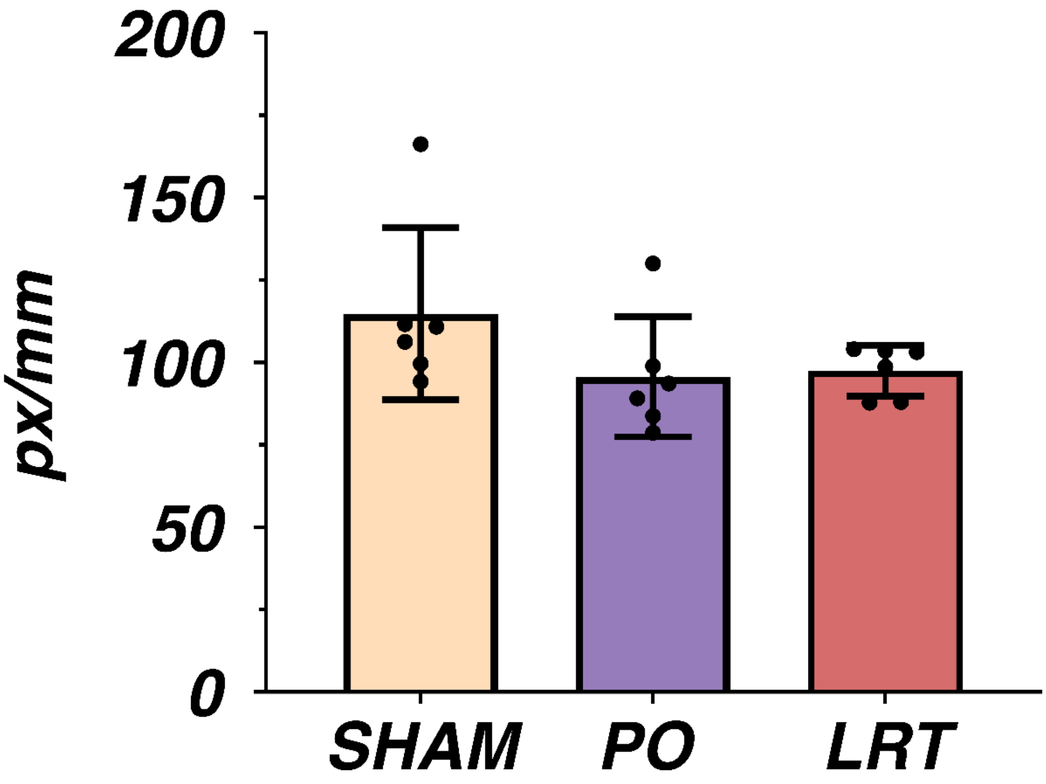


**Figure S6. Digital image correlation (DIC) spatial resolution (in pixels per millimeter) for sham, PO, and LRT groups. Bars represent mean ± SD. No significant differences were detected between groups using a Kruskal-Wallis test (sham: 114.8 ± 26.1 px/mm; PO: 95.6 ± 18.3 px/mm; LRT: 97.5 ± 7.7 px/mm).**

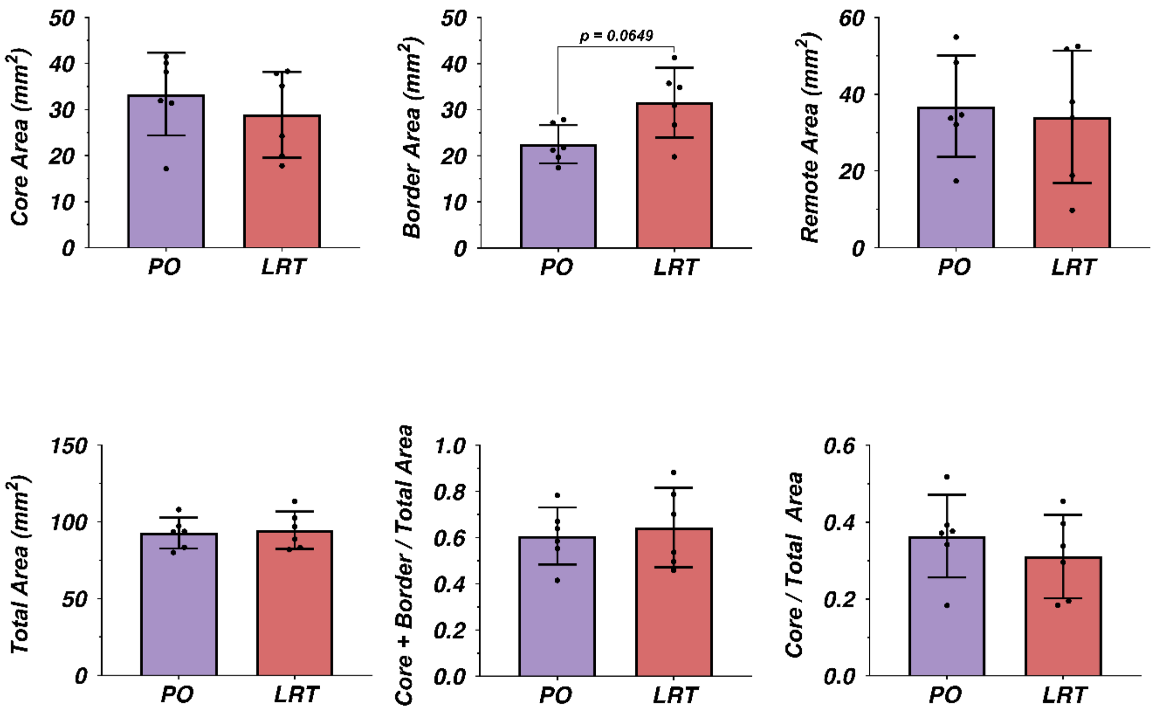


**Figure S7. Quantification of infarcted sample regions determined via a k-means clustering algorithm: infarct core area (A), infarct border area (B; *p* < 0.1 using a Mann-Whitney U-test), remote area (C), total sample area (D), total infarct area (core + border) normalized to total sample area (E), and infarct core area normalized to total sample area (F). Each bar represents the mean ± SD across all PO and LRT samples.**

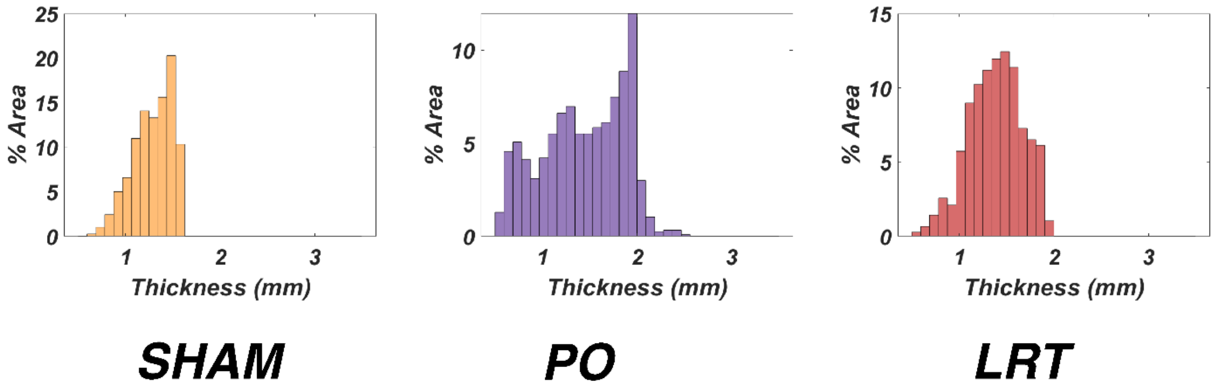


**Figure S8. Percent surface area at each thickness value for the representative sham, PO, and LRT samples shown in Figure 3A and throughout the main manuscript.**

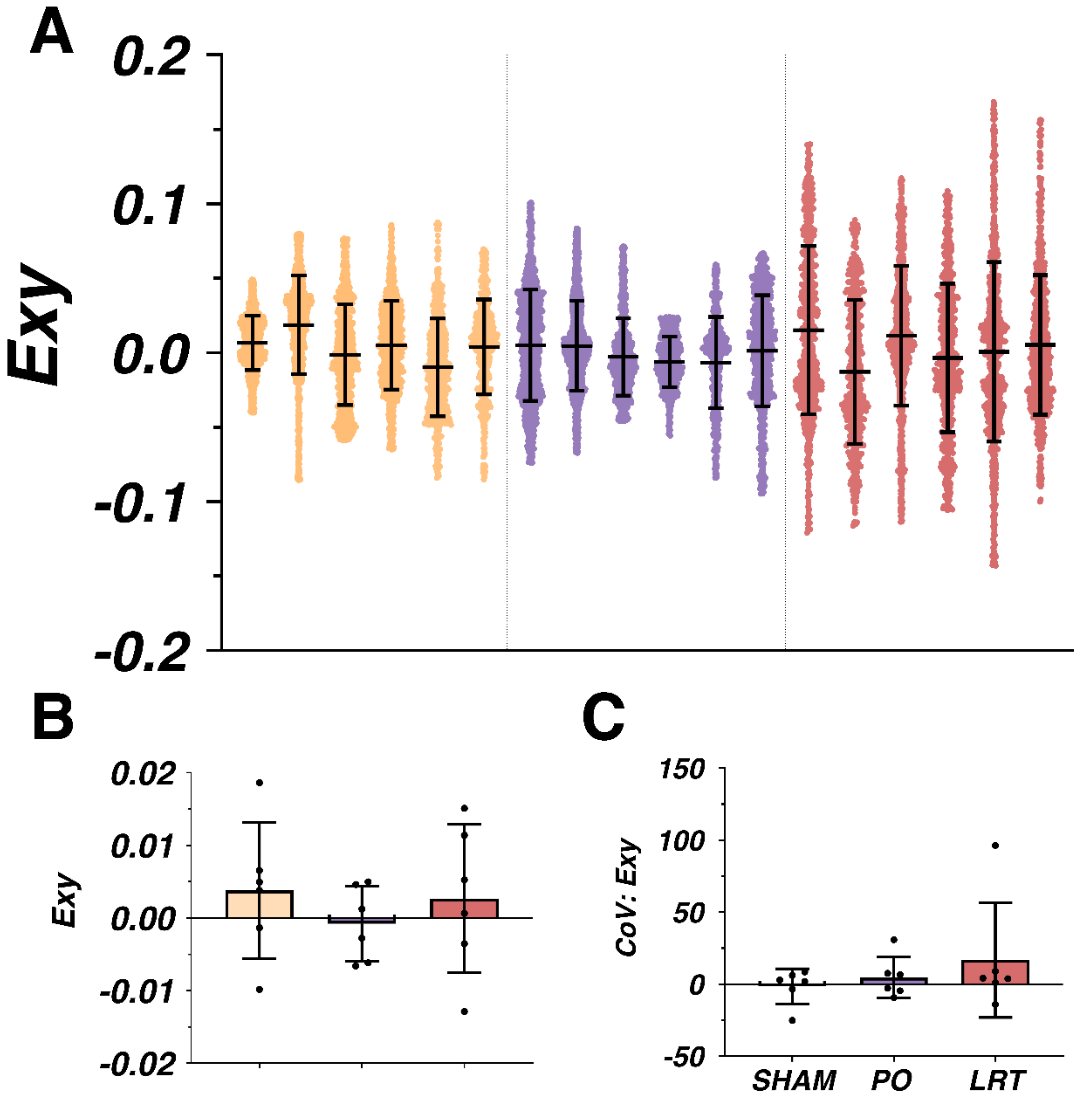


**Figure S9. (A) Scatter plots showing full-field shear strain (*Exy*) distributions at peak equibiaxial extension for all samples tested. Bar plots comparing mean *Exy* (B) and the coefficients of variation (CoV) in *Exy* (C) at peak equibiaxial extension. No inter-group statistically significant differences were detected via a Mann-Whitney *U*-test.**

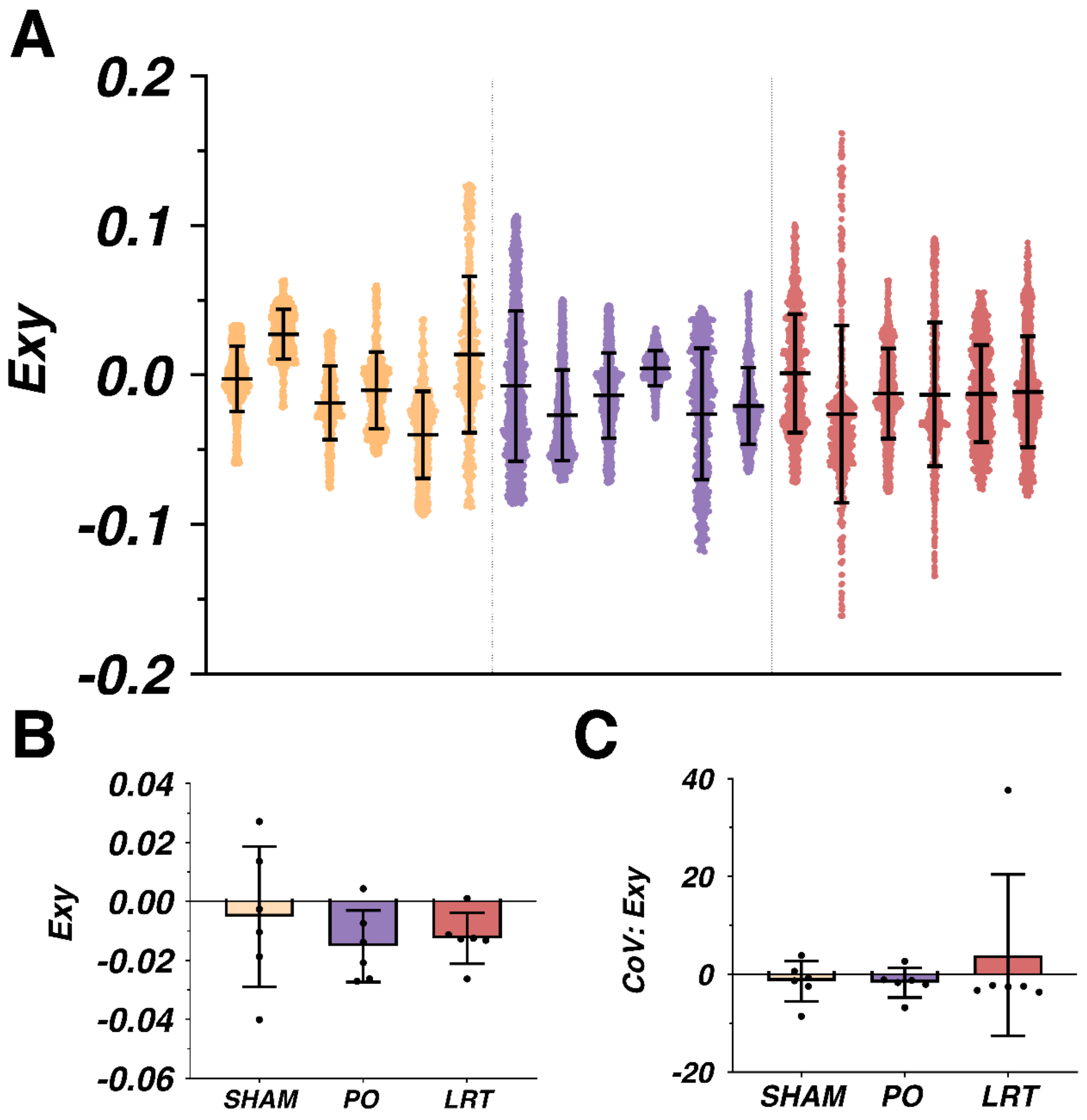


**Figure S10. (A) Scatter plots showing full-field shear strain (*Exy*) distributions at peak Strip X extension for all samples tested. Bar plots comparing mean *Exy* (B*)* and the coefficients of variation (CoV) in *Exy* (C) at peak Strip X extension. No inter-group statistically significant differences were detected via a Mann-Whitney *U*-test.**

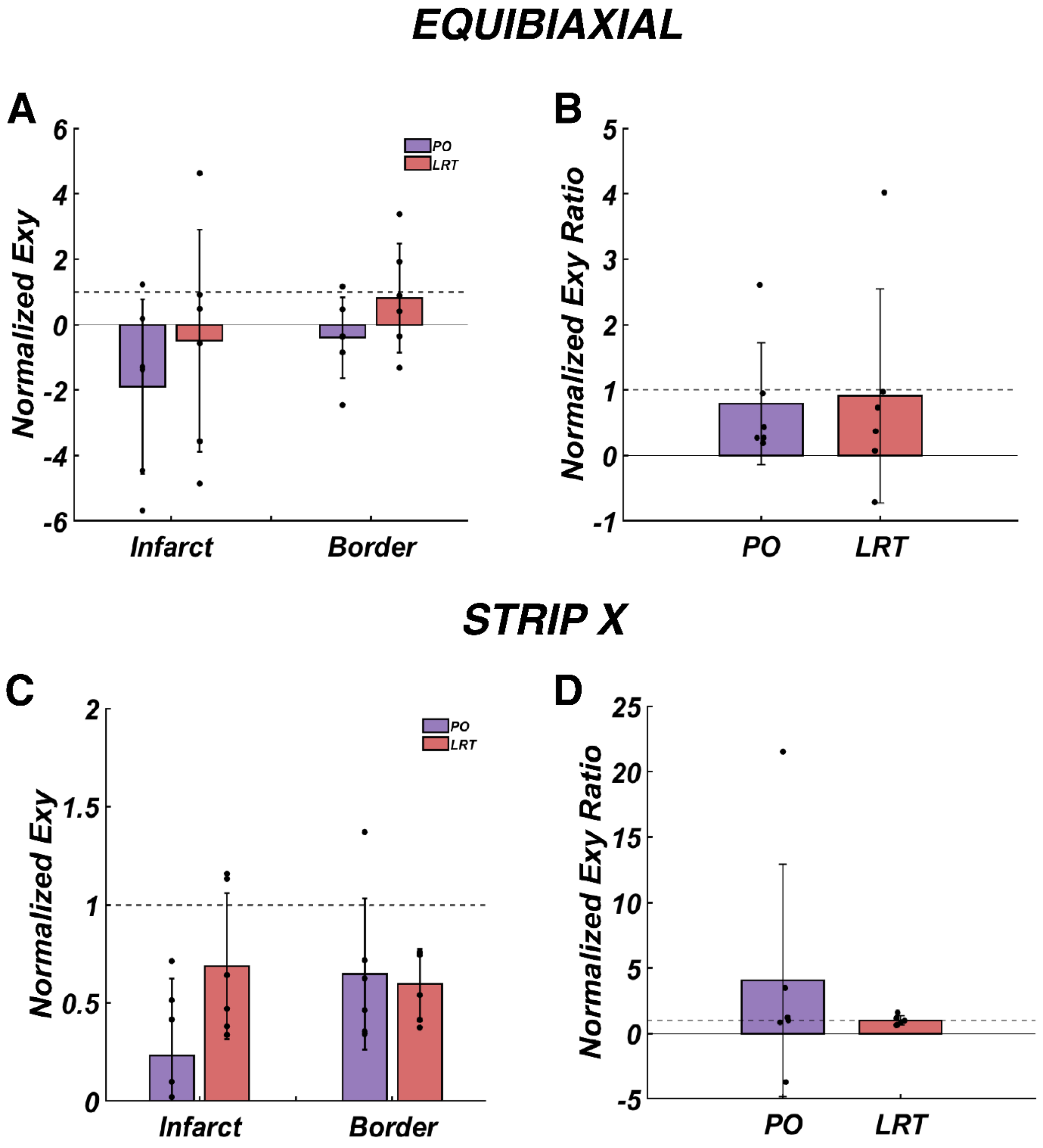


**Figure S11. Normalized infarct core and border *Exy* (A) and the border-to-core *Exy* ratios (B) at peak equibiaxial extension. Normalized infarct core and border Exy (C) and the border-to-core *Exy* ratios (D) at peak Strip X extension. No statistically significant differences were detected when regional values were compared to a remote value of one using a Wilcoxon signed-rank test.**

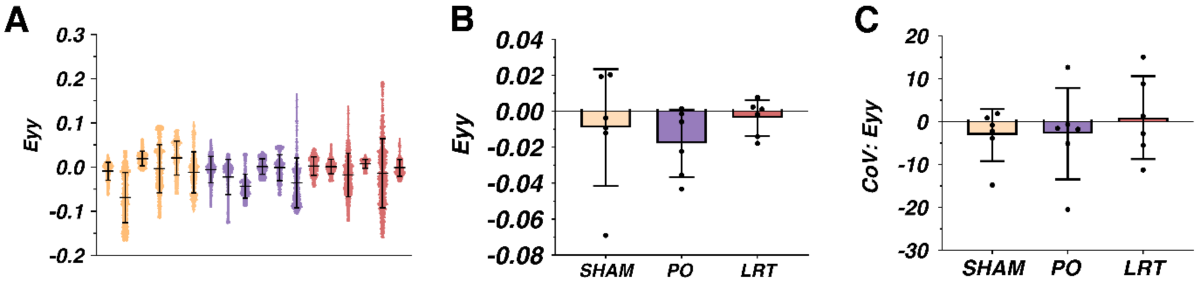


**Figure S12. (A) Scatter plots showing full-field vertical strain (*Eyy*) distributions at peak Strip X extension for all samples tested. Bar plots comparing mean *Eyy* (B), and the CoV in *Eyy* (C) at peak Strip X extension. No inter-group statistically significant differences were detected via a Mann-Whitney *U*-test.**

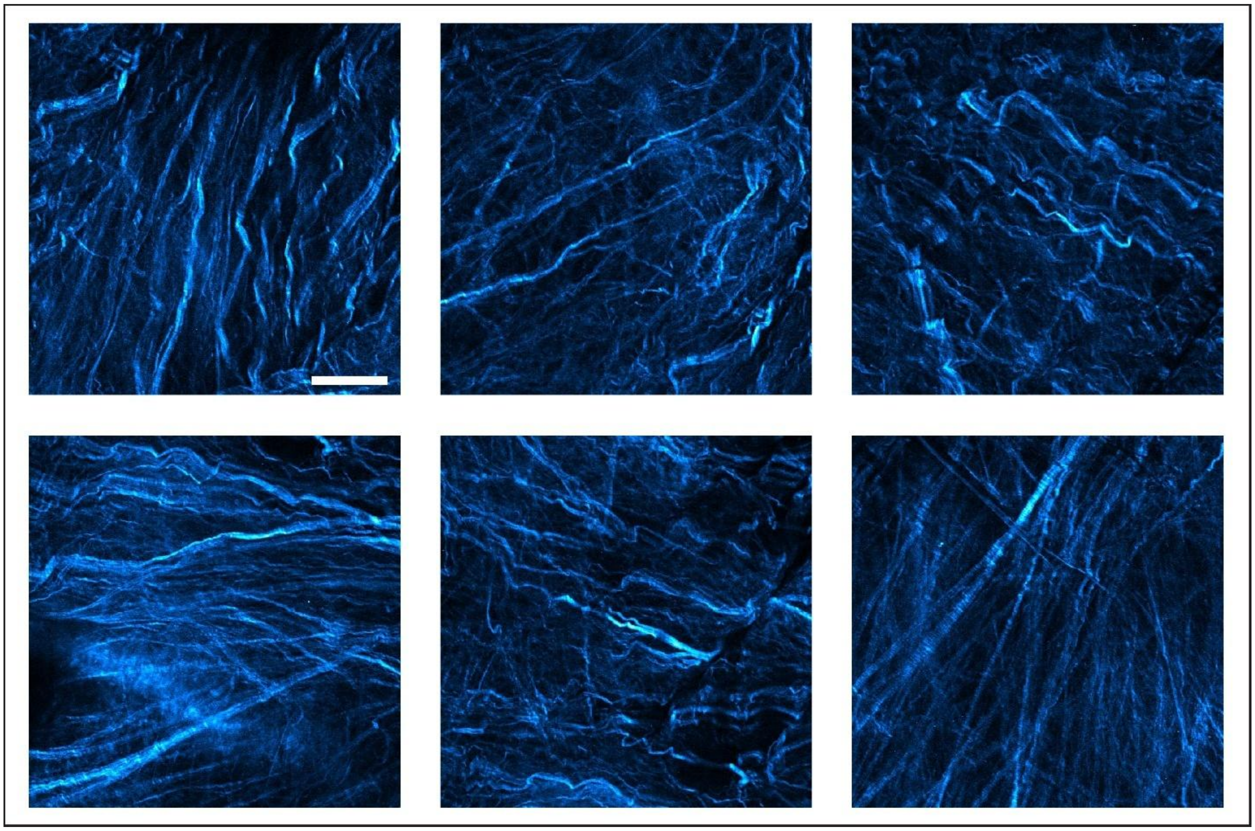

**Figure S13. SHG images of collagen fibers from six randomly selected locations within a decellularized sham LV sample. The scale bar indicates 70 microns.**

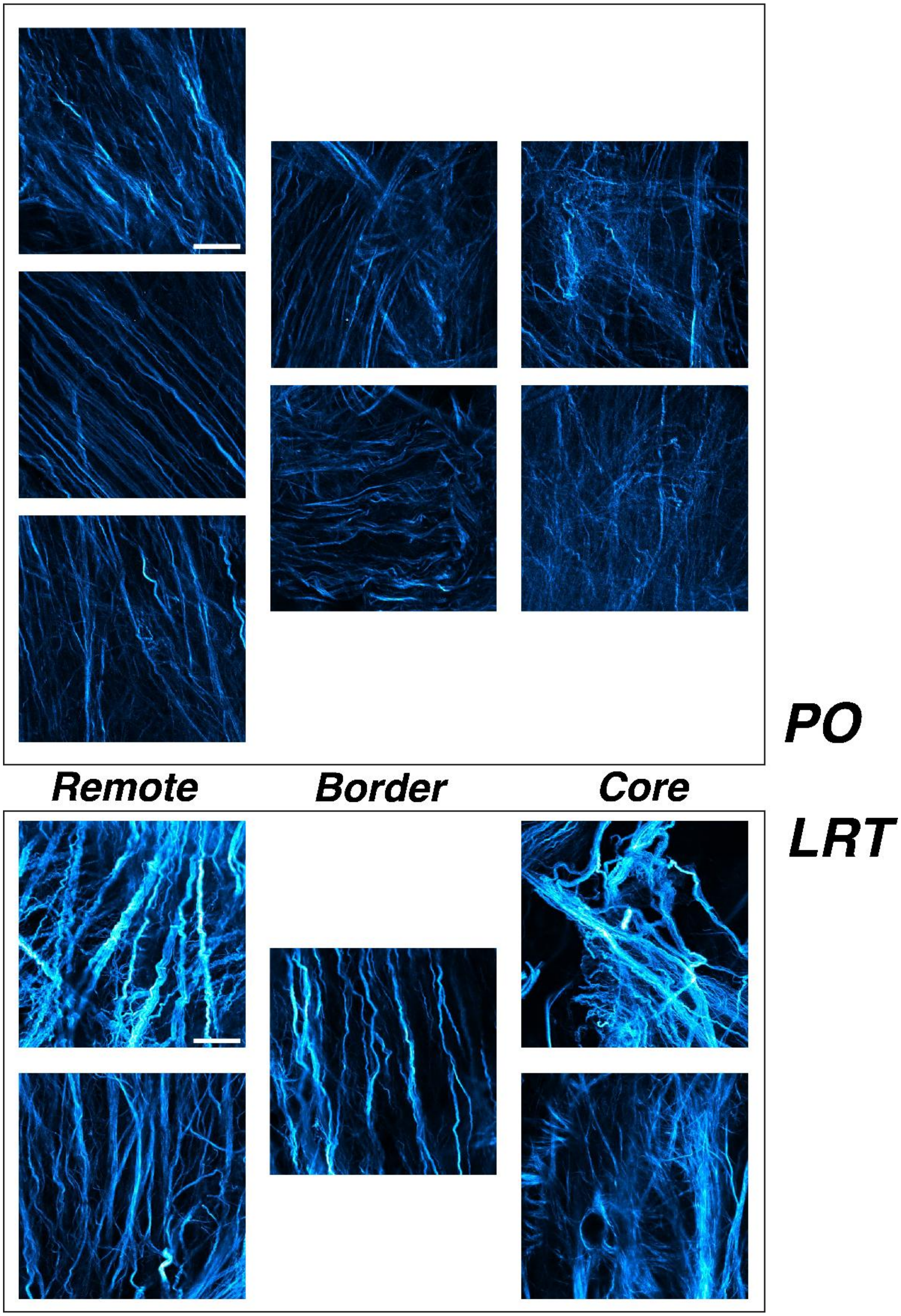


**Figure S14. Representative SHG images of collagen fibers from the infarct core, border, and remote LV regions in a decellularized PO and LRT sample. Scale bars indicate 70 microns.**

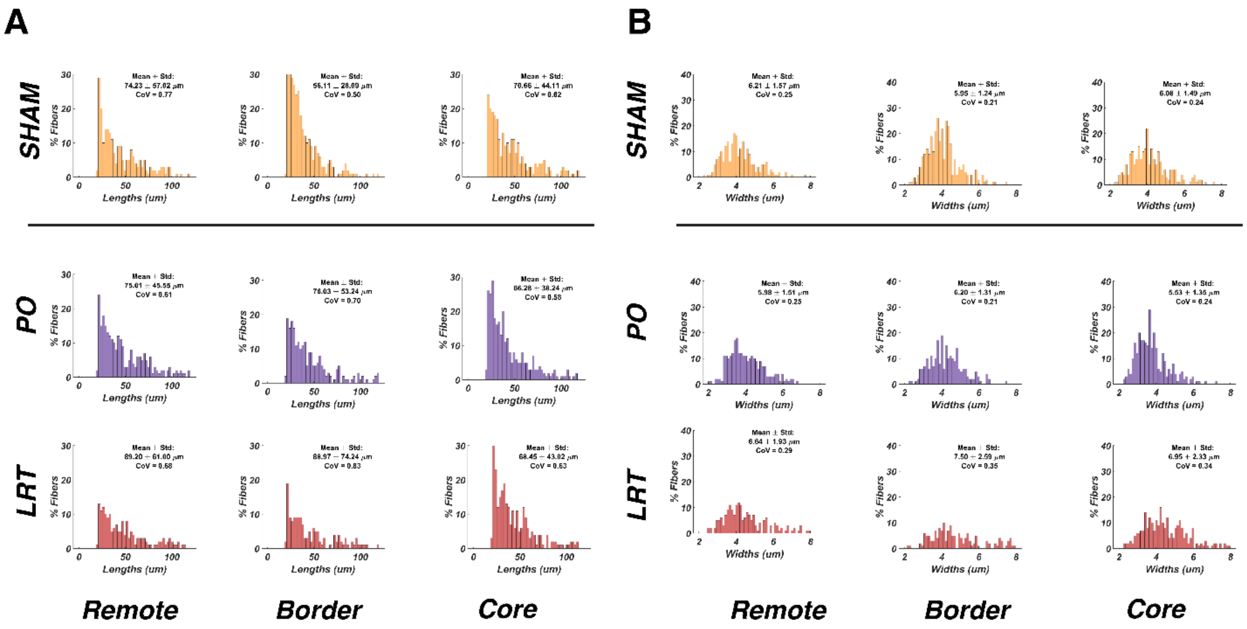


**Figure S15. Quantification of collagen fiber length (A; in um) and width (B; in um) from CT-Fire in SHG images shown in Figure 8A.**

**Table S1. P-values from Mann-Whitney U-test comparisons of PO and LRT groups unloaded and loaded regional areas as well as the change in these areas during each extension. Note the unloaded areas are the same for every extension.**

| | Unloaded Area P-Value | | | Loaded Area P-Value | | | Delta Area P-Value | |
|---|---|---|---|---|---|---|---|---|
| Extension | Infarct Core | Infarct Border | Extension | Infarct Core | Infarct Border | Extension | Infarct Core | Infarct Border |
| 1 | 0.4848 | 0.0649 | 1 | 0.3939 | 0.026 | 1 | 0.5887 | 0.026 |
| 2 | 0.4848 | 0.0649 | 2 | 0.3939 | 0.0411 | 2 | 0.6991 | 0.026 |
| 3 | 0.4848 | 0.0649 | 3 | 0.3939 | 0.0649 | 3 | 0.5887 | 0.0411 |
| 4 | 0.4848 | 0.0649 | 4 | 0.3939 | 0.0649 | 4 | 0.8182 | 0.026 |
| 5 | 0.4848 | 0.0649 | 5 | 0.3939 | 0.026 | 5 | 0.8182 | 0.0152 |
| 6 | 0.4848 | 0.0649 | 6 | 0.3939 | 0.0649 | 6 | 0.6991 | 0.0411 |
| 7 | 0.4848 | 0.0649 | 7 | 0.3939 | 0.0649 | 7 | 0.1797 | 0.0087 |
| 8 | 0.4848 | 0.0649 | 8 | 0.4848 | 0.0931 | 8 | 0.8182 | 0.2403 |
| 9 | 0.4848 | 0.0649 | 9 | 0.3939 | 0.0411 | 9 | 0.6991 | 0.0022 |
| 10 | 0.4848 | 0.0649 | 10 | 0.3939 | 0.0411 | 10 | 0.9372 | 0.0087 |
| 11 | 0.4848 | 0.0649 | 11 | 0.5887 | 0.0649 | 11 | 0.9999 | 0.026 |
| 12 | 0.4848 | 0.0649 | 12 | 0.3939 | 0.0152 | 12 | 0.5887 | 0.0087 |
| 13 | 0.4848 | 0.0649 | 13 | 0.3939 | 0.0649 | 13 | 0.9372 | 0.0152 |
| 14 | 0.4848 | 0.0649 | 14 | 0.3939 | 0.0649 | 14 | 0.5887 | 0.0411 |
| 15 | 0.4848 | 0.0649 | 15 | 0.3939 | 0.026 | 15 | 0.6991 | 0.0087 |

**Table S2. P-values from Mann-Whitney U-test comparisons of the unloaded and loaded border-to-core area ratios as well as the change in these area ratios during each extension between PO and LRT groups. Note the unloaded ratios are the same for every extension.**

| **Extension** | **Unloaded Ratio P-Value** | **Loaded Ratio P-Value** | **Delta Ratio P-Value** |
|---|---|---|---|
| 1 | 0.0411 | 0.0260 | 0.00216 |
| 2 | 0.0411 | 0.0411 | 0.0260 |
| 3 | 0.0411 | 0.0411 | 0.0649 |
| 4 | 0.0411 | 0.0411 | 0.0411 |
| 5 | 0.0411 | 0.0411 | 0.0152 |
| 6 | 0.0411 | 0.0411 | 0.310 |
| 7 | 0.0411 | 0.0411 | 0.00866 |
| 8 | 0.0411 | 0.0411 | 0.394 |
| 9 | 0.0411 | 0.0411 | 0.0152 |
| 10 | 0.0411 | 0.0411 | 0.0152 |
| 11 | 0.0411 | 0.0411 | 0.180 |
| 12 | 0.0411 | 0.0411 | 0.00866 |
| 13 | 0.0411 | 0.0411 | 0.0152 |
| 14 | 0.0411 | 0.0411 | 0.0649 |
| 15 | 0.0411 | 0.0411 | 0.0152 |

**Table S3. P-values from Mann-Whitney U-test comparisons of unloaded and loaded border width as well as the change in this width during each extension between PO and LRT groups. Note the unloaded widths are the same for every extension.**

| **Extension** | **Unloaded Border Width P-Value** | **Loaded Border Width P-Value** | **Delta Border Width P-Value** |
|---|---|---|---|
| 1 | 0.818 | 0.699 | 0.0260 |
| 2 | 0.818 | 0.818 | 0.0931 |
| 3 | 0.818 | 0.699 | 0.310 |
| 4 | 0.818 | 0.818 | 0.310 |
| 5 | 0.818 | 0.699 | 0.310 |
| 6 | 0.818 | 0.818 | 0.485 |
| 7 | 0.818 | 0.699 | 0.240 |
| 8 | 0.818 | 0.699 | 0.240 |
| 9 | 0.818 | 0.818 | 0.132 |
| 10 | 0.818 | 0.699 | 0.394 |
| 11 | 0.818 | 0.589 | 0.310 |
| 12 | 0.818 | 0.818 | 0.0649 |
| 13 | 0.818 | 0.818 | 0.180 |
| 14 | 0.818 | 0.818 | 0.132 |
| 15 | 0.818 | 0.699 | 0.240 |
